# Hemispheric asymmetries in resting-state EEG and fMRI are related to approach and avoidance behaviour, but not to eating behaviour or BMI

**DOI:** 10.1101/692012

**Authors:** Filip Morys, Lieneke Janssen, Elena Cesnaite, Frauke Beyer, Isabel Garcia-Garcia, Jana Kube, Deniz Kumral, Franziskus Liem, Nora Mehl, Keyvan Mahjoory, Anne Schrimpf, Michael Gaebler, Daniel Margulies, Arno Villringer, Jane Neumann, Vadim Nikulin, Annette Horstmann

**Affiliations:** Leipzig University Medical Centre, IFB Adiposity Diseases, 04103 Leipzig, Germany; Department of Neurology, Max Planck Institute for Human Cognitive and Brain Sciences, 04103 Leipzig, Germany; Montreal Neurological Institute, McGill University, Montreal, QC, Canada; Subproject A1/A5, CRC1052 “Obesity Mechanisms”, University of Leipzig, Leipzig, Germany; Brandenburg University of Technology Cottbus-Senftenberg, 03013 Cottbus, Germany; MindBrainBody Institute at the Berlin School of Mind and Brain, Humboldt-Universitaet zu Berlin, Berlin, Germany; University Research Priority Program “Dynamics of Healthy Aging”, University of Zurich, CH-8006 Zurich, Switzerland; Max Planck Research Group for Neuroanatomy & Connectivity, Max Planck Institute for Human Cognitive and Brain Sciences, Leipzig, Germany; Faculty of Psychology, Technical University Dresden, 01062 Dresden, Germany; University of Muenster, Institute for Biomagnetism and Biosignal Analysis, 48149 Muenster, Germany; Brain and Spine Institute, 75013 Paris, France; Ernst-Abbe-Hochschule – University of Applied Sciences, 07745 Jena, Germany; Centre for Cognition and Decision Making, Institute for Cognitive Neuroscience, National Research University Higher School of Economics, Moscow, Russia; Department of Neurology, Charité – Medical University Berlin, 10117 Berlin, Germany; Department of Psychology and Logopedics, Faculty of Medicine, University of Helsinki

**Author notes:** **Corresponding author:** Filip Morys, Montreal Neurological Institute, McGill University, 3801 Rue Université, Montréal, QC H3A 2B4, Canada. **Data availability** The data that support the findings of this study are available from the corresponding author upon reasonable request.

**Keywords:** Approach/avoidance behaviour, resting-state, EEG, fMRI, hemispheric asymmetries, obesity, BMI

## Abstract

Much of our behaviour is driven by two motivational dimensions – approach and avoidance. These have been related to frontal hemispheric asymmetries in clinical and resting-state EEG studies: approach was linked to higher activity of the left relative to the right hemisphere, while avoidance was related to the opposite pattern. Increased approach behaviour, specifically towards unhealthy foods, is also observed in obesity and has been linked to asymmetry in the framework of the right-brain hypothesis of obesity. Here, we aimed to replicate previous EEG findings of hemispheric asymmetries for self-reported approach/avoidance behaviour and to relate them to eating behaviour. Further, we assessed whether resting fMRI hemispheric asymmetries can be detected and whether they are related to approach/avoidance, eating behaviour, and BMI. We analysed 3 samples: Sample 1 (n=117) containing EEG and fMRI data from lean participants, and Samples 2 (n=89) and 3 (n=152) containing fMRI data from lean, overweight, and obese participants. While in Sample 1 approach in women was related to EEG and fMRI hemispheric asymmetries, in Samples 2 and 3 this effect was not significant. Here, hemispheric asymmetries were neither related to BMI nor eating behaviour. Our study partly replicates previous EEG findings regarding hemispheric asymmetries and indicates that this relationship could also be captured using fMRI. Our findings suggest that eating behaviour and obesity are likely to be mediated by mechanisms not directly relating to frontal asymmetries in neuronal activation quantified with EEG and fMRI.

## 1 Introduction

A sizeable proportion of our everyday actions is driven by approach (e.g., reaching for a tasty biscuit) and avoidance (e.g., running away from a big spider) tendencies. Such tendencies can be considered fundamental motivational dimensions that steer (not only) human behaviour (Davidson & Hugdahl, 1995). These two dimensions at the core of the framework of behavioural inhibition and activation systems (BIS and BAS, respectively; Gray, 1981; Gray & McNaughton, 1992) and can, for example, be assessed by means of the self-report BIS/BAS questionnaire (Carver & White, 1994). Literature on individual differences in terms of inhibition and activation systems is broad and mostly focuses on disorders such as depression, anxiety, substance addictions or obesity (Dietrich, Federbusch, Grellmann, Villringer, & Horstmann, 2014; Johnson, Turner, & Iwata, 2003; Morgan et al., 2009). There is experimental evidence that both substance addictions and obesity are related to increased approach behaviour towards problematic stimuli: while substance abuse relates to approach towards cigarettes, marijuana or alcohol substances, obesity relates to approach tendencies towards unhealthy food cues (Cousijn et al., 2012; Mehl, Morys, Villringer, & Horstmann, 2019; Mehl, Mueller-Wieland, Mathar, & Horstmann, 2018; Wiers et al., 2013; Wiers et al., 2014). Furthermore, obesity and higher body mass index (BMI) were shown to relate to BIS/BAS scores in a gender-dependent fashion, with positive correlations in women, and negative correlations in men (Dietrich et al., 2014).

Regarding the neural correlates of approach/avoidance behaviours, literature suggests differential engagement of left and right frontal brain areas, such as the Brodmann area 9 or 10, and reward-related regions of the brain, such as the nucleus accumbens or the ventral tegmental area (Aberg, Doell, & Schwartz, 2015; Tomer et al., 2013). The left hemisphere is more strongly engaged in approach, and the right in avoidance behaviours (Aberg et al., 2015; Davidson, 1993, 1994; Sutton & Davidson, 1997; Tomer, Goldstein, Wang, Wong, & Volkow, 2008). A seminal study showed that higher alpha power, which is believed to represent inhibitory control (Bazanova & Vernon, 2014; Klimesch, Sauseng, & Hanslmayr, 2007), in right frontal brain areas (relative to the left) measured in resting-state EEG (rsEEG), was associated with increased approach behaviour (Sutton & Davidson, 1997). This was explained by downregulated right hemispheric activity, since alpha power has previously been linked to cortical inhibition by top-down control and suppression of task-irrelevant brain regions (Bazanova, 2012; Klimesch et al., 2007). A number of studies showed similar functional asymmetries in reward regions such as the ventral tegmental area and nucleus accumbens using positron emission tomography (Tomer et al., 2013) and task-based fMRI (Aberg et al., 2015) during reward and punishment learning. These findings suggest that hemispheric asymmetries and their relationship to approach/avoidance behaviours can be quantified using a range of neuroimaging tools. However, the relationship between approach/avoidance behaviours and hemispheric asymmetries in resting-state fMRI (rsfMRI) has not yet been investigated.

Since obesity is related to altered approach/avoidance behaviours, it might also be related to hemispheric asymmetries. This hypothesis is grounded in the right-brain theory of obesity, which posits that hypoactivation of the right prefrontal cortex is an underlying factor of obesity (Alonso-Alonso & Pascual-Leone, 2007). It is based on findings of increased eating behaviour after damages to right-hemispheric anterior brain areas (Regard & Landis, 1997; Short, Broderick, Patton, Arvanitakis, & Graff-Radford, 2005). It is also supported by EEG experiments showing a higher right hemispheric bias for restrained eaters, a predominantly inhibitory feature (Silva, Pizzagalli, Larson, Jackson, & Davidson, 2002) and a positive relationship of left hemispheric bias with disinhibition and hunger (Ochner, Green, van Steenburgh, Kounios, & Lowe, 2009) as measured with the three factor eating questionnaire (TFEQ; Stunkard & Messick, 1985). The above-mentioned studies, however, did not investigate a direct link between obesity measures, such as BMI and hemispheric asymmetries. Furthermore, due to the method of choice (EEG), those studies could focus mainly on cortical brain structures. Since obesity is often related to functional alterations in dopaminergic subcortical structures (Cone, Chartoff, Potter, Ebner, & Roitman, 2013; Friend et al., 2016; Geiger et al., 2009; Horstmann, Fenske, & Hankir, 2015; Narayanaswami, Thompson, Cassis, Bardo, & Dwoskin, 2013; Stice, Yokum, Burger, Epstein, & Small, 2011; Volkow, Wang, Fowler, & Telang, 2008; Vucetic, Carlin, Totoki, & Reyes, 2012), focusing on subcortical asymmetries using suitable neuroimaging techniques, such as fMRI, might further our knowledge regarding the neural correlates of obesity.

In this study, we addressed three aims using three independent samples. First, we aimed to conceptually replicate the previous findings from the literature concerning hemispheric asymmetries in terms of EEG alpha power, self-reported approach/avoidance (BIS/BAS), and eating behaviour (TFEQ, cognitive control and disinhibition) questionnaires. This was done in a large sample of predominantly lean participants (Sample 1, 117 participants). Second, we aimed to show that the relationship of approach/avoidance, eating behaviour and rsEEG asymmetry can be extended to rsfMRI in the same sample. Here, we also aimed to investigate hemispheric asymmetries in subcortical structures, which cannot be easily done using EEG. Third, we aimed to establish the existence of obesity-related hemispheric asymmetries in rsfMRI by investigating self-reported eating behaviours (TFEQ), approach/avoidance behaviours (BIS/BAS), and BMI in two samples including lean, overweight, and obese participants (Sample 2, 89 participants; Sample 3, 152 participants). The three samples enabled us to provide a conceptual replication of previous studies, while at the same time expanding existing knowledge to new behavioural measures and methods.

We hypothesised that higher self-reported approach behaviour (BAS) would be related to increased left vs. right hemispheric activity, whereas higher self-reported avoidance (BIS) would be related to increased right vs. left hemispheric activity in both rsEEG and rsfMRI. Furthermore, increased cognitive control of food intake was expected to be related to higher right vs. left hemispheric activity, whereas higher disinhibition was expected to be related to increased left vs. right hemispheric activity. Lastly, we hypothesised higher BMI to be related to increased left vs. right hemispheric activity. We further aimed to investigate whether approach/avoidance-related hemispheric asymmetries can be measured using both EEG and fMRI neuroimaging, as was previously done in a different context, e.g. language research (Mazza & Pagano, 2017; Powell et al., 2006).

## 2 Materials and methods

Analysed data were parts of different projects, all of which were conducted according to the Declaration of Helsinki and approved by local Ethics Committees (University of Leipzig, Germany – Sample 1 and 2; Montclair State University and Nathan Kline Institute – Sample 3). All participants gave their written informed consent prior to participation.

### 2.1 Participants

#### 2.1.1 Sample 1

Sample 1 consisted of 117 healthy, right-handed, predominantly lean participants aged 20-35 years (mean age: 25 years, mean BMI: 23.01 kg/m^2^, range: 17.95-37.80 kg/m^2^; 42 women, Table S1) taken from the ‘Leipzig Study for Mind-Body-Emotion Interactions’ (Babayan et al., 2019). Exclusion criteria included: history of psychiatric or neurological disease, substance abuse, hypertension, MRI-related contraindications (cf. Table 1 in (Babayan et al., 2019). Data available for this sample included self-reported eating (TFEQ) and approach/avoidance behaviour (BIS/BAS) questionnaires, anthropometric data (BMI), rsEEG and rsfMRI (Table S2). For analysis of EEG data, 1 participant was excluded due to an unresponsive electrode of interest, which resulted in a sample of 116 participants. For analysis of fMRI data, 3 participants were excluded due to data pre-processing problems (failed registration), and 3 additional participants were excluded due to excessive head motion during data acquisition (criterion: maximum framewise displacement exceeding 2.3mm; Power, Barnes, Snyder, Schlaggar, & Petersen, 2012), which resulted in a sample of 111 participants.

**Table 1.** Results of multiple regression analyses investigating the relationship between EEG asymmetry indices and approach/avoidance questionnaire measures. Statistically significant coefficients have been marked in bold. Note that the p-value threshold after Bonferroni correction for four separate regression analyses is 0.0125. RC – rotated component, RR – reward responsivity

|  | Frontal |  |  |  | Parietal |  |  |  |
| --- | --- | --- | --- | --- | --- | --- | --- | --- |
|  | Alpha |  | Low alpha |  | Alpha |  | Low alpha |  |
|  | Beta | p-value | Beta | p-value | Beta | p-value | Beta | p-value |
| BAS fun | -0.14 | 0.5275 | 0.14 | 0.2508 | 0.16 | 0.4318 | 0.27 | 1.0000 |
| BAS fun * gender | -0.02 | 1.0000 | -0.13 | 0.6545 | -0.29 | 0.3721 | -0.27 | 1.0000 |
| BAS drive | 0.32 | 0.3420 | **0.66** | **0.0108\*** | 0.31 | 0.5217 | 0.08 | 0.8630 |
| BAS drive * gender | -0.51 | 0.1000 | **-0.88** | **0.0018\*** | -0.34 | 0.4815 | 0.06 | 0.7450 |
| BAS RR | 0.54 | 0.0477 | 0.34 | 0.1083 | 0.08 | 0.6545 | -0.22 | 0.3270 |
| BAS RR * gender | -0.54 | 0.0660 | -0.35 | 0.1747 | -0.22 | 0.4184 | 0.03 | 0.6860 |
| BAS - BIS | -0.38 | 0.2312 | -0.35 | 0.6863 | -0.77 | 0.0304 | -0.53 | 0.1350 |
| BAS – BIS * gender | 0.57 | 0.1294 | 0.59 | 0.1241 | **0.97** | **0.0078\*** | 0.56 | 0.4130 |
| Age | 0.02 | 0.8039 | 0.04 | 0.7255 | 0.12 | 0.3122 | 0.00 | 1.0000 |
| BMI | **0.31** | **0.0024\*** | **0.25** | **0.0076\*** | -0.13 | 0.1793 | -0.11 | 0.2530 |
| Gender | 0.45 | 0.0330 | 0.44 | 0.0373 | 0.42 | 0.4214 | 0.27 | 0.2350 |

#### 2.1.2 Sample 2

Sample 2 consisted of 89 healthy, right-handed, lean, overweight and obese participants aged 20-37 years (mean age: 27 years, mean BMI: 29.54 kg/m^2^, range: 17.67-59.78 kg/m^2^; 73 women, Table S1). The data were collected at the Max Planck Institute for Human Cognitive and Brain Sciences in Leipzig. This sample was created by merging data of two different studies from our lab investigating decision-making in obesity. Subsample 1 consisted of 56 lean, overweight and obese women, whereas Subsample 2 consisted of 33 participants with obesity, men and women (Mehl et al., 2019). Data available for both subsamples were self-reported eating (TFEQ) and approach/avoidance behaviour (BIS/BAS) questionnaires, anthropometric data (BMI), and rsfMRI data (Table S2). Exclusion criteria were: history of psychiatric or neurological disease, substance abuse, hypertension, MRI-related contraindications. No participants had to be excluded during data analysis.

#### 2.1.3 Sample 3

Sample 3 consisted of participants from an open database of the enhanced Nathan Kline Institute-Rockland Sample (NKI; http://fcon_1000.projects.nitrc.org/indi/enhanced/; releases up to 6^th^). From this database, we selected rsfMRI data of 152 healthy, right-handed lean, overweight and obese participants aged 18-35 years (mean age: 24 years, mean BMI: 26.40 kg/m^2^, range: 16.26-49.96 kg/m^2^; 84 women, Table S1) with Beck Depression Inventory scores below 18 indicating lack of depressive symptoms (Beck, Steer, Ball, & Ranieri, 1996). Additional data available for this sample were self-reported eating behaviour (TFEQ) data and anthropometric data (BMI; Table S2).

### 2.2 Questionnaire data

To investigate how hemispheric asymmetries reflect approach and avoidance behaviours, we used the BIS/BAS (behavioural inhibition system / behavioural activation system) questionnaire (Carver & White, 1994). This questionnaire was administered Samples 1 and 2. It consists of 4 different scales: three subscales reflecting BAS (drive, reward responsivity and fun seeking) and a subscale reflecting BIS. According to Carver and White, the drive scale reflects persistent pursuit of desired goals; the reward responsivity scale focuses on positive responses to rewarding events; the fun seeking scale reflects a desire for new rewards and the inclination to approach a rewarding event. The BIS scale, on the other hand, describes individual sensitivity to punishment.

With regard to the self-reported eating behaviour, we used the three factor eating questionnaire (TFEQ; Stunkard & Messick, 1985). It describes eating behaviour on three dimensions: cognitive control for food (CC), disinhibition (DI), and susceptibility to hunger (H). In this study, we were predominantly interested in the first two factors, as they might reflect avoidance and approach behaviour towards food, respectively.

### 2.3 Neuroimaging data

#### 2.3.1 EEG data acquisition – Sample 1

In this study, participants completed three assessment sessions in three days (Babayan et al., 2019). The first assessment day included a cognitive test battery and a set of questionnaires. On the second assessment day, rsEEG data were acquired, which consisted of 16 blocks, each lasting 1 min of intermittent eyes closed (EC) and eyes open (EO) conditions, summing up to a total duration of 8 min per condition. RsEEG was recorded in an acoustically shielded room with 62 active electrodes (Brain Vision ActiCAP; Brain Products GmbH, Munich, Germany) placed according to the international standard 10–20 extended localization system, also known as 10-10 system (Oostenveld & Praamstra, 2001), all referenced to FCz electrode, with the ground electrode placed on the sternum. Electrooculographic (EOG) activity was recorded with one electrode placed below the right eye. EEG signals were sampled at 2500 Hz and band-pass filtered between 0.015 Hz and 1 kHz, the amplifier was set to 0.1 µV amplitude resolution, and electrode impedance was kept below 5kΩ.

#### 2.3.2 fMRI data acquisition – Sample 1

For Sample 1, MRI data were collected with a 3T Siemens Verio scanner (Siemens, Erlangen, Germany). We analysed T2*-weighted rsfMRI, MP2RAGE and fieldmap data. RsfMRI data parameters: 657 volumes, TE=30ms, FA=69°, TR=1400ms, 64 slices in an interleaved order, voxel size: 2.3×2.3×2.3mm^3^, FoV: 202mm, multiband acceleration factor: 4, acquisition time: 15 minutes. MP2RAGE parameters: TE=2.92ms, FA1=4°, FA2=5°, TR=2500ms, TI1=700ms, TI2=2500ms, voxel size: 1×1×1mm^3^, FoV: 256mm.

#### 2.3.3 fMRI data acquisition – Sample 2

MRI data for both of the subsamples of this sample were collected with a 3T Siemens Skyra scanner. We analysed T2*-weighted rsfMRI, MPRAGE and fieldmap data. RsfMRI parameters: 320 volumes, TE=22ms, FA=90°, TR=2000ms, 40 slices in an ascending order, voxel size: 3.0×3.0×2.5mm^3^, FoV: 192mm, acquisition time: 11 minutes. MPRAGE parameters: TE=2.01ms, FA=9°, TR=2300ms, TI=900ms, voxel size: 1×1×1mm^3^, FoV: 256mm.

#### 2.3.4 fMRI data acquisition – Sample 3

For Sample 3, MRI data were collected with a 3T Siemens Trio scanner. We analysed T2*-weighted rsfMRI and MPRAGE data. RsfMRI parameters (http://fcon_1000.projects.nitrc.org/indi/enhanced/mri_protocol.html): 900 volumes in an interleaved order, TE=30ms, FA=60°, TR=645ms, 40 slices, voxel size: 3.0×3.0×2.5mm^3^, FoV: 222mm, multiband acceleration factor: 4, acquisition time: 10 minutes. MPRAGE parameters: TE=2.52ms, FA=9°, TR=2600ms, TI=900ms, voxel size: 1×1×1mm^3^, FoV: 256mm.

### 2.4 Data pre-processing

#### 2.4.1 EEG data - Sample 1

EEG data were pre-processed using EEGLAB toolbox (version 14.1.1b; Delorme & Makeig, 2004) and custom Matlab (MathWorks, Inc, Natick, Massachusetts, USA) scripts. EEG time series were band-pass filtered between 1-45 Hz (4th order back and forth Butterworth filter) and downsampled to 250 Hz. EC and EO segments were extracted and concatenated which resulted in an 8-min block per condition. Artifactual channels and time segments were removed after visual inspection. Principal component analysis (PCA) was performed to reduce the data dimensionality to N components (N≥30) that explained 95% of the total variance. A pre-processing with PCA was used for the following independent component analysis (Infomax; Bell & Sejnowski, 1995) was used to reject components related to eye movements, muscle activity, and heartbeats. For further analyses, the pre-processed EEG time series were transformed to the common average reference.

#### 2.4.2 fMRI data – Samples 1 and 2

fMRI data pre-processing for Samples 1 and 2 was identical and was done within the Nipype framework (Gorgolewski et al., 2011). In short, the pre-processing steps included discarding the first five functional volumes, motion correction (FSL MCFLIRT; Jenkinson, Bannister, Brady, & Smith, 2002), distortion correction (FSL FUGUE; Jenkinson, Beckmann, Behrens, Woolrich, & Smith, 2012), coregistration of the temporal mean image to the individual’s anatomical image (bbregister; Greve & Fischl, 2009), denoising (rapidart and aCompCor; Behzadi, Restom, Liau, & Liu, 2007), spatial normalisation to MNI 152 2mm (Sample 1), and 3mm (Sample 2) standard space (ANTs; Avants et al., 2011). The details of the pipeline are described in Mendes et al., 2017.

#### 2.4.3 fMRI data – Sample 3

fMRI data pre-processing for Sample 3 data was also done within the Nipype framework (Gorgolewski et al., 2011). In short, the pre-processing steps included discarding first five functional volumes, motion correction (FSL MCFLIRT; Jenkinson et al., 2002), denoising (rapidart and aCompCor; Behzadi et al., 2007), removal of linear and quadratic signal trends), spatial normalisation to a 3mm standard MNI 152 space (FSL FNIRT; Jenkinson et al., 2012). The details of the pipeline are described in Liem et al., 2017. Note that the bandpass filtering described in Liem et al. was not performed for our data, since further statistical analysis of the fMRI data (fALFF) require them to be unfiltered.

### 2.5 Neuroimaging measures

#### 2.5.1 Aim 1: EEG replication analysis

In this step we, attempted to directly replicate previous findings from Sutton and Davidson (1997) showing a positive correlation of left hemispheric bias with BAS – BIS differential scores. BAS – BIS differential scores represent a balance between the two systems and higher scores represent the dominance of the BAS system. In this first analysis and in this analysis only, we calculated an absolute EEG asymmetry index in frontal areas by subtracting absolute alpha power (8-12Hz) in the F3 electrode (left) from absolute alpha power in the F4 electrode (right; asymmetry index: R-L) for mean values of EO and EC conditions together.

We then wanted to extend previous findings concerning EEG hemispheric bias and approach/avoidance behaviour to eating behaviour (as measured by the TFEQ). As rsfMRI was collected with eyes open to prevent subjects from falling asleep, our main analysis focused on EEG data from the eyes open condition in order to compare it with fMRI findings. We additionally conducted EEG analyses with relative alpha power of eyes open and eyes closed conditions to investigate whether potential effects observed only in the eyes open condition are specific to this condition or can be extended to other conditions as well. The mean of eyes open and eyes closed was also used to make the analysis closer to previous EEG studies on hemispheric bias, which also used this measure.

We focused on alpha power in the broader spectrum (8-12Hz) and in the narrower spectrum for low alpha (8-10Hz) for our analysis. While the broader alpha frequency band (8–12 Hz) has been previously linked to cortical inhibition by top-down control (Bazanova, 2012; Klimesch et al., 2007), low alpha power (8-10 Hz) was previously shown to reflect general attentional demands, basic alertness, vigilance, and arousal (Klimesch et al., 2007; Petsche, Kaplan, von Stein, & Filz, 1997). Including both of the measures allowed us to replicate previous results obtained using broadband alpha, and confine possible mechanistic interpretations to, for example, general attentional demands (by using low alpha). For this analysis, as opposed to the direct replication described in the previous paragraph, we used relative alpha power to control for inter-individual differences in contaminating factors like skull thickness or other properties of the scalp and meninges that might affect tissue conductivity and influence electrical signal captured at the sensor level (Babiloni et al., 2011). Relative power in broadband alpha and low alpha frequency ranges were calculated by firstly taking the mean of the squared amplitude obtained after filtering the signal (4th order back and forth Butterworth filter) in the 8-12Hz and the 8-10Hz frequency ranges, respectively, and then dividing it by the power within the frequency range of 4-40Hz. In line with Sutton and Davidson (Sutton & Davidson, 1997), relative alpha power measures were calculated in the pair of frontal electrodes F4 and F3. We also included a parietal pair, P4 and P3, as a control to investigate whether the observed relationship with frontal asymmetries was topographically specific.

Previous research on hemispheric asymmetries used an absolute asymmetry index (Sutton & Davidson, 1997), while in our study we calculated a relative asymmetry index using the following equation: (R-L)/(R+L). By accounting for inter-individual differences in alpha power magnitude, these relative indices capture asymmetries better than the absolute R-L difference and increase interpretability (Hiroshige & Dorokhov, 1997; Pivik et al., 1993). After calculation of asymmetry indices, we excluded outliers from all variables of interest using the *a priori* defined criterion (see section 2.6).

#### 2.5.2 Aim 2+3: Hemispheric asymmetries in fMRI

After pre-processing (sections 2.4.2 and 2.4.3), analysis of fMRI data in all 3 samples was identical. To be able to conceptually compare EEG results with fMRI results, the fractional amplitude of low-frequency fluctuations (fALFF) was used as a measure of resting-state brain activity (Zou et al., 2008). fALFF is usually defined as the ratio of power in the frequency range of 0.01-0.1Hz and the power within the entire detectable frequency range. However, the samples had different sampling frequencies during fMRI data collection (i.e. repetition time, TR) and thus different detectable frequency ranges. To be able to better compare results between the samples, the denominator of the fALFF ratio was fixed to 0.00Hz – 0.25Hz, reflecting the frequency range for the sample with the highest TR. This analysis was performed in the Nipype framework using CPAC (Configurable Pipeline for the Analysis of Connectomes, version 1.0.3, https://fcp-indi.github.io/) f/ALFF function. To compare EEG and fMRI results from our original analysis, we defined a set of regions of interest (ROI) for the fMRI analysis. Based on previous literature (Giacometti, Perdue, & Diamond, 2014; Herwig, Satrapi, & Schönfeldt-Lecuona, 2003; Towle et al., 1993), we determined 7 ROIs that corresponded to brain areas measured by the EEG analysis in frontal (F3/F4) and parietal (P3/P4) electrodes: Brodmann areas 8, 9, 10, 46 reflecting frontal contributions, Brodmann area 7, postcentral gyrus, and paracentral gyrus reflecting parietal contributions. These ROIs were defined using pickatlas (Maldjian, Laurienti, Kraft, & Burdette, 2003). Since fMRI allows to investigate subcortical brain areas, for which hemispheric asymmetries have been shown (Aberg et al., 2015; Mathar et al., 2017; Tomer et al., 2008), we additionally tested a hemispheric bias in the ventral tegmental area (sphere with a 6mm radius, coordinates based on (Aberg et al., 2015; Adcock, Thangavel, Whitfield-Gabrieli, Knutson, & Gabrieli, 2006); L: x=−4, y=−15, z=−9; R: x=5, y=−14, z=−8), and the nucleus accumbens (sphere with a 6mm radius, coordinates based on (Aberg et al., 2015; Neto, Oliveira, Correia, & Ferreira, 2008); L: x=−9, y=9, z=− 8; R: x=9, y=8, z=−8). For each ROI, which was defined separately for the left and for the right hemisphere, we extracted mean fALFF using SPM 12 (Wellcome Department of Cognitive Neurology, London, United Kingdom). A relative asymmetry index was calculated as follows: (L-R)/(L+R). Note that this is an inverse index compared to the one we used for EEG data, since we hypothesised that measures used in EEG and fMRI analysis are inversely correlated, due to physiological the phenomena that they are thought to measure (i.e. inhibition vs. activation, respectively). This let us directly compare relationships of EEG and fMRI data with behavioural measures, which was one of the aims of the study.

### 2.6 Statistical analysis

For each of the variables of interest, outliers were excluded based on an *a priori* criterion: 2.2*interquartile range below or above the first or third quartile, respectively (Hoaglin & Iglewicz, 1987; Hoaglin, Iglewicz, & Tukey, 1986; Tukey, 1977). Further, all regression p-values were corrected for multiple comparisons using Bonferroni correction, that is, by dividing the alpha value 0.05 by the number of regressions performed on the same dataset. All statistical analyses were performed using R (version 3.2.3) within JupyterNotebook.

#### 2.6.1 Aim 1: EEG replication analysis

To directly replicate Sutton’s and Davidson’s research (1997), for each participant we calculated the differential BAS – BIS score. We then removed outliers from both measures of interest (rsEEG and questionnaire data) and correlated BAS – BIS scores with absolute alpha asymmetry indices. To analyse the data, we performed Pearson’s correlation of the obtained EEG asymmetry indices (section 2.5.1) and the BAS – BIS scores. Final sample size for this analysis after outlier exclusion was 109 participants.

To investigate the relationship between approach and avoidance behaviours and hemispheric bias as a direct replication of previous studies, we performed four separate multiple regression analyses with asymmetry indices from relative frontal alpha power, relative parietal alpha power, relative frontal low alpha power, and relative parietal low alpha power as outcome variables. This was done separately for the EO condition, and for the mean of EO and EC conditions. Predictors included BAS fun, BAS drive, and BAS reward responsivity as well as BAS – BIS scores. Note, however, that due to the relatively low BMI range in Sample 1, in the analysis of this sample BMI served as a variable of no interest. To investigate whether gender influences the relationship between questionnaire measures and hemispheric bias, we added an interaction term with gender for each of the questionnaire variables. This was done because previous findings show that approach/avoidance behaviours might be gender dependent (Dietrich et al., 2014). To control for age differences, we also added this information to the model as a predictor. This and all following regression analyses were calculated using permutation tests in the ‘lmPerm’ R package (Bonferroni corrected α=0.0125).

To analyse self-reported eating behaviour, similar regression analyses were performed as described in the previous paragraph with different questionnaire variables: cognitive control and disinhibition (TFEQ) and their interactions with gender, and BMI, and age as variables of no interest (Bonferroni corrected α=0.0125).

#### 2.6.2 Aim 2: EEG-fMRI correspondence

##### 2.6.2.1 Correlations between EEG and fMRI

First, we wanted to directly investigate the relationship of EEG asymmetries (frontal and parietal) and whole-brain fALFF asymmetries in Sample 1 to investigate the relationships between EEG and fMRI measures. Whole-brain fALFF asymmetries were calculated by means of 1) flipping left and right hemispheres in fALFF images (left becomes right and *vice versa*), 2) subtracting the flipped image from the original image, 3) adding the flipped image to the original image, and 4) dividing the image obtained in step 2 by the image obtained in step 3. This resulted in an image of voxel-wise values corresponding to the asymmetry index (L-R)/(L+R) (on the left side of the image, and (R-L)/(R+L) index of the right side of the brain image). A significant correlation between the EEG asymmetry index as calculated in 2.5.1 and whole-brain fALFF asymmetries would indicate that those two measures, even though methodologically very distinct, measure similar brain processes. This analysis was performed in SPM12 using a general linear model with voxel-wise fALFF asymmetries as an outcome variable and the EEG asymmetry index as an explanatory variable. Results were thresholded on a voxel-level with a 0.001 threshold, and corrected for multiple comparisons using the whole-brain 0.05 FWE-corrected threshold.

##### 2.6.2.2 Relationships between fMRI hemispheric asymmetries and approach/avoidance and eating behaviours in Sample 1

To investigate relationships of fMRI hemispheric bias with approach/avoidance and eating behaviours, we first used rotated principal component analysis (PCA) on the ROI imaging data (asymmetry indices calculated for mean fALFF values per ROI). This was done to reduce the number of comparisons in further analyses (Jolliffe & Cadima, 2016). We used the varimax rotation, which drives component loadings (correlations of components and original variables) either towards zero or towards a maximum possible value, decreasing a number of components with medium loadings, which are difficult to interpret (Jolliffe, 2002; M. B. Richman, 1986; M. L. B. Richman, 1987). As a criterion for retaining components we chose the minimum cumulative variance explained to be over 70% (Jolliffe, 2002). This resulted in 5 components for each of the samples.

Further, to investigate relationships of fMRI hemispheric bias and approach/avoidance behaviour, we performed a similar analysis to the one using EEG data. Five rotated principal components were defined as outcome measures, and predictors included BAS fun, BAS drive, BAS reward responsivity as well as BIS - BAS scores and their interaction with gender. Additionally, we included BMI and age as variables of no interest (Bonferroni corrected α=0.01).

A similar analysis was performed to investigate relationships between fMRI hemispheric bias and eating behaviour. It included similar predictors as the EEG investigation of eating behaviour – cognitive control and disinhibition and their interaction with gender. Outcome variables were 5 rotated principal components. We added BMI and age as variables of no interest (Bonferroni corrected α=0.01).

#### 2.6.3 Aim 3: fMRI investigations in samples including participants with obesity - relationship of hemispheric bias and self-reported behaviours

Investigations of approach/avoidance behaviours in Sample 2 were performed similarly to the ones in Sample 1. Five rotated components were defined as outcome variables, and predictors included BIS/BAS questionnaire measures, their interaction with gender, and BMI. Age was added as a regressor of no interest (Bonferroni corrected α=0.0100).

A similar analysis was performed to investigate associations of self-reported eating behaviour and hemispheric asymmetries for Samples 2 and 3. Predictor variables included eating questionnaire measures and their interaction with gender, BMI, age (regressor of no interest), while outcome variables were 5 rotated components (Bonferroni corrected α=0.0100).

### 2.7 Final sample sizes

Approach/avoidance behaviours: For the direct replication EEG analysis, the sample size was 113 after outlier exclusion. For measurement of approach/avoidance behaviours within Sample 1 (eyes open), to be able to compare EEG results with fMRI findings, sample size was reduced to 100 participants due to outlier exclusion in EEG and in fMRI data matrices simultaneously. The same analysis for eyes open and eyes closed was done on a sample of 93 participants. Sample 2 after outlier exclusion consisted of 87 participants.

Eating behaviours: Final size of Sample 1 was 95 for eyes open condition, and 88 for mean of eyes open and eyes closed conditions. Sample 2 consisted of 87 participants, and the size of Sample 3 was 138 participants. We decided to exclude outliers separately for analysis of BIS/BAS data and TFEQ data to maximise sample sizes between analyses which were anyway performed as separate multiple regressions. Hence, a potential participant who would be excluded as an outlier in the BIS/BAS data analysis could be retained for the TFEQ data analysis.

## 3 Results

### 3.1 Aim 1: EEG replication analysis – Sample 1

In this analysis, we aimed to directly replicate findings of Sutton and Davidson (1997) of increased hemispheric bias (R-L; F4 – F3 electrodes, absolute alpha power, mean values for EO and EC conditions) being related to increased BAS – BIS differential scores. We did not find a significant relationship between those variables (r_(113)_=0.121, p=0.202). Partial correlation after controlling for BMI, age, and gender also did not reveal a significant relationship (r_(113)_=0.094, p=0.325).

Next, we attempted to expand previous findings linking EEG and approach/avoidance behaviours to 1) additional spectra to improve specificity and interpretability of findings, 2) additional questionnaire measures to improve specificity of the findings. We therefore investigated relationships between EEG parietal and frontal asymmetry indices as measured by the relative broad alpha power, as used by Sutton & Davis, and relative low alpha power. In addition to the standard broad alpha power spectrum used in previous studies, low alpha power spectrum due to its specific physiological meaning (general attentional demands, basic alertness, vigilance, and arousal; Klimesch et al., 2007; Petsche et al., 1997) allowed us to more precisely interpret relationships between hemispheric asymmetries and behaviour. Here, we used the improved, relative asymmetry index: (R-L)/(R+L). For questionnaire data we included BAS fun seeking, drive, reward responsivity in addition to BAS – BIS differential scores. Firstly, we investigated the eyes open condition. Results of this analysis (Table 1) indicate a significant relationship of BAS drive and frontal hemispheric bias in low alpha frequency for women only (BAS drive: p=0.0108, BAS drive*gender: p=0.0018). This is shown by an interaction of BAS drive with gender, and a significant main effect of BAS drive. In this analysis, women were coded as 0 and were the reference category, hence the main effect of BAS drive shows that this relationship is true for women, because in this case all other interaction terms including gender are also equal to zero. A similar relationship was not significant for broad alpha power. For scatter plots of these relationships see Figure 1. We also observed a significant interaction of BAS – BIS scores and gender for alpha parietal asymmetry indices, suggesting gender to influence the relationship between BAS – BIS and EEG asymmetries. However, we do not interpret significant findings for BMI, age, and gender, since those variables were added to the model as covariates of no interest. In the analysis of mean of eyes open and eyes closed conditions we found no significant effects (Table S3). This suggests that the asymmetry findings are specific to the eyes open condition only.

**Figure 1.**
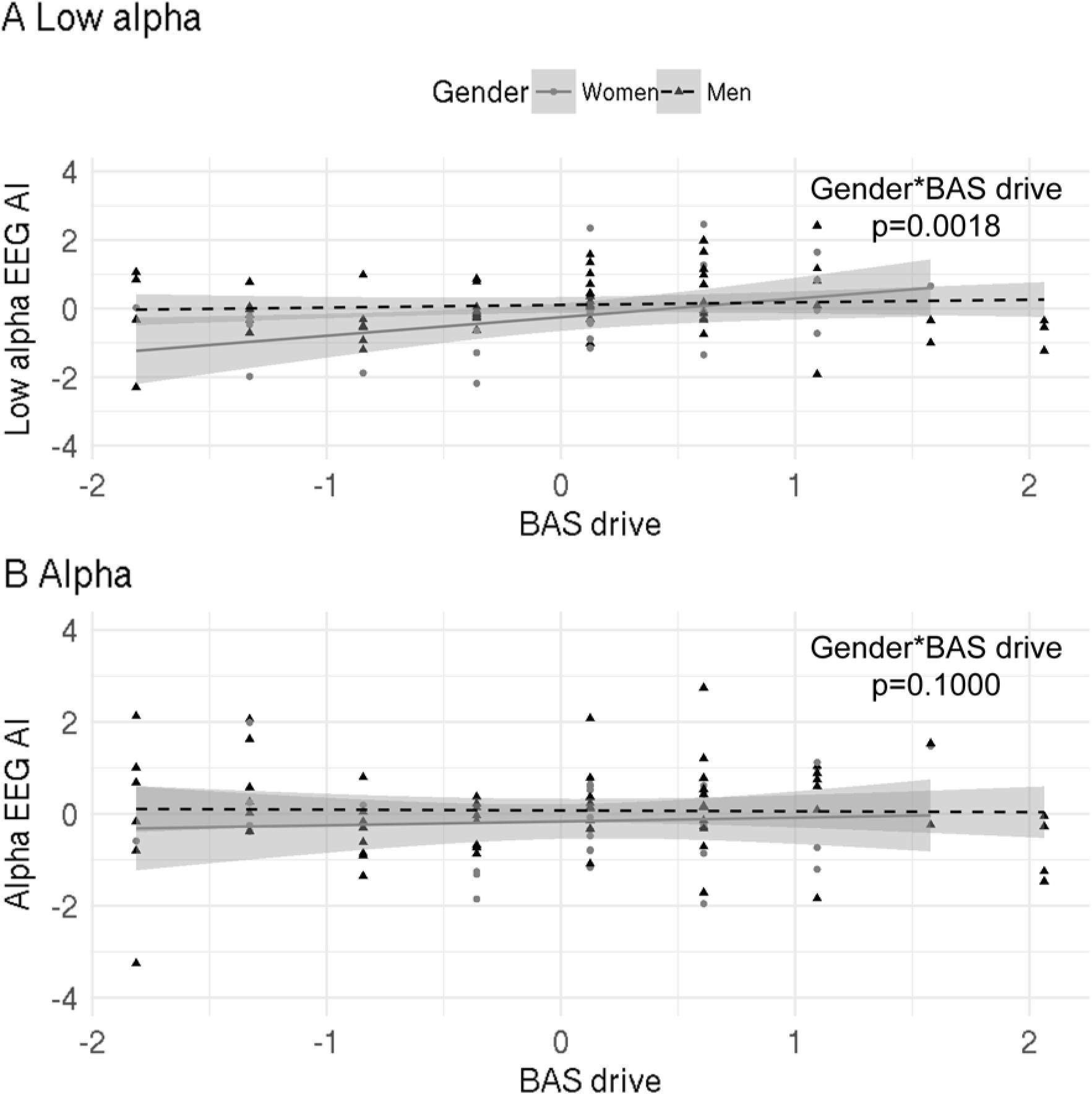
Relationship between low/full alpha EEG asymmetry index (AI) and BAS drive scores. Index used: (R-L)/(R+L). Triangles/dots represent data points, dashed/bold lines represent the best fit, and grey shaded areas are 95% confidence intervals. **A** – significant correlation of hemispheric asymmetries and behavioural measures in the low alpha spectrum (beta: −0.88, p=0.0018); **B** – not significant correlation of hemispheric asymmetries and behavioural measures in the broad alpha spectrum showing that the asymmetries are specific to the low alpha spectrum (beta: 0.51, p=0.1000). AI – asymmetry index, R – right, L – left.

Further, we investigated the relationship between the TFEQ and the EEG hemispheric bias. Predictor variables in this case included cognitive control, disinhibition, and their interactions with gender (BMI and age entered as regressors of no interest). Here, we did not find any significant associations for eyes open condition, or for mean of eyes open and eyes closed conditions. Detailed results of these analyses can be found in Tables S4 and S5.

### 3.2 Aim 2: fMRI correspondence analysis – Sample 1

Firstly, we investigated direct relationships between EEG asymmetries (using the relative asymmetry index (R-L)/(R+L)) and whole-brain fALFF asymmetry measures in the same sample. This analysis did not produce significant results (Table 2), suggesting no correspondence between rsEEG and rsfMRI hemispheric bias measures.

**Table 2.**
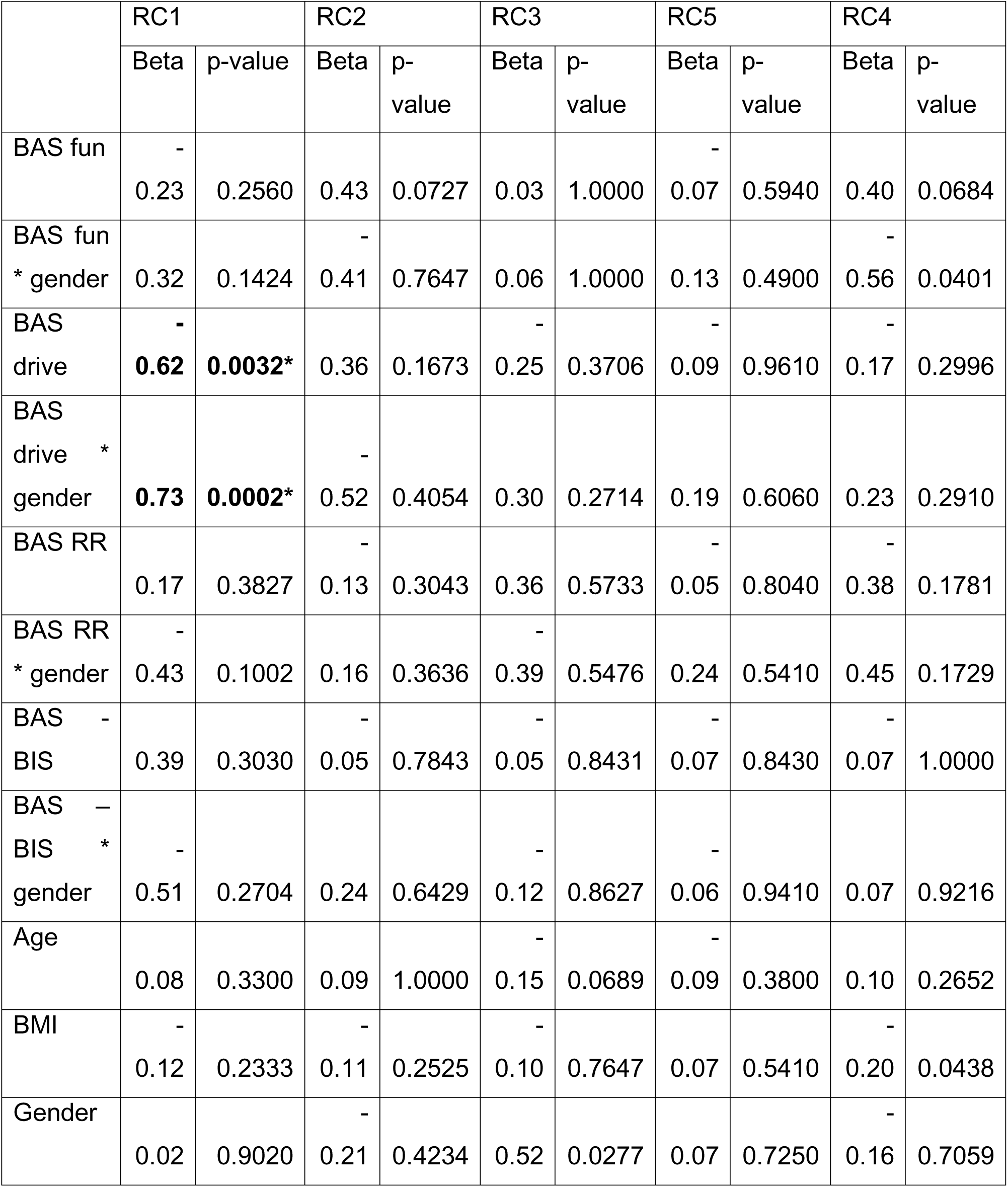
Results of multiple regression analyses investigating the relationship between fMRI asymmetry indices (Sample 1) and approach/avoidance questionnaire measures. Note that the p-value threshold after Bonferroni correction for five separate regression analyses is 0.0100. The components have been ordered according to decreasing variance explained (Table 3). RC – rotated component, RR – reward responsivity

Next, we investigated relationships between fMRI relative asymmetry indices (L-R)/(L+R) and approach/avoidance behaviours in Sample 1. The analysis included 5 retained components describing asymmetry data and questionnaire variables – BAS fun, BAS drive, BAS reward responsivity, BAS – BIS and their interactions with gender. Additionally, we included BMI and age as covariates of no interest.

We found a significant interaction effect of BAS Drive and gender on the rotated component 1 (RC1), and a main effect of BAS drive on RC1 with contributions from the BA9, BA8, and ventral tegmental area (p=0.0032 and p=0.0002, respectively). Results of this analysis can be found in Table 2. For a visualisation of the data see Figure 2. Loading of each of the RCs in the PCA analysis can be found in Table 3. It indicates that the RC1 was mostly influenced by the BA9, BA8, and VTA. For visualisation purposes we present raw (before PCA) ROI data (BA9, BA8, VTA) relationships with BAS Drive scores for men and women in Figure 3.

**Table 3.** Component loadings and cumulative variance explained for each of the rotated components (RC, Sample 1) in the BIS/BAS analysis. ROIs represent 9 regions of interest selected for the fMRI analyses. BA - Brodmann area; NAcc - nucleus accumbens; VTA – ventral tegmental area; ParaCG – paracentral gyrus; PostcG – postcentral gyrus; ROI - region of interest.

| ROI | RC1 | RC2 | RC3 | RC5 | RC4 |
| --- | --- | --- | --- | --- | --- |
| BA10 | 0.08 | 0.68 | 0.16 | 0.41 | -0.22 |
| BA9 | 0.73 | 0.38 | -0.11 | -0.04 | -0.13 |
| BA8 | 0.71 | 0.27 | 0.10 | 0.14 | -0.17 |
| BA46 | 0.24 | 0.75 | -0.02 | -0.12 | 0.10 |
| NAcc | -0.01 | -0.02 | 0.93 | -0.05 | 0.01 |
| VTA | 0.79 | -0.24 | 0.00 | 0.18 | 0.18 |
| BA7 | 0.17 | 0.04 | -0.09 | 0.92 | 0.04 |
| ParacG | -0.06 | 0.67 | -0.49 | 0.06 | -0.01 |
| PostcG | -0.06 | 0.00 | 0.01 | 0.02 | 0.96 |
| Cumulative variance explained | 0.20 | 0.39 | 0.52 | 0.64 | 0.76 |

**Figure 2.**
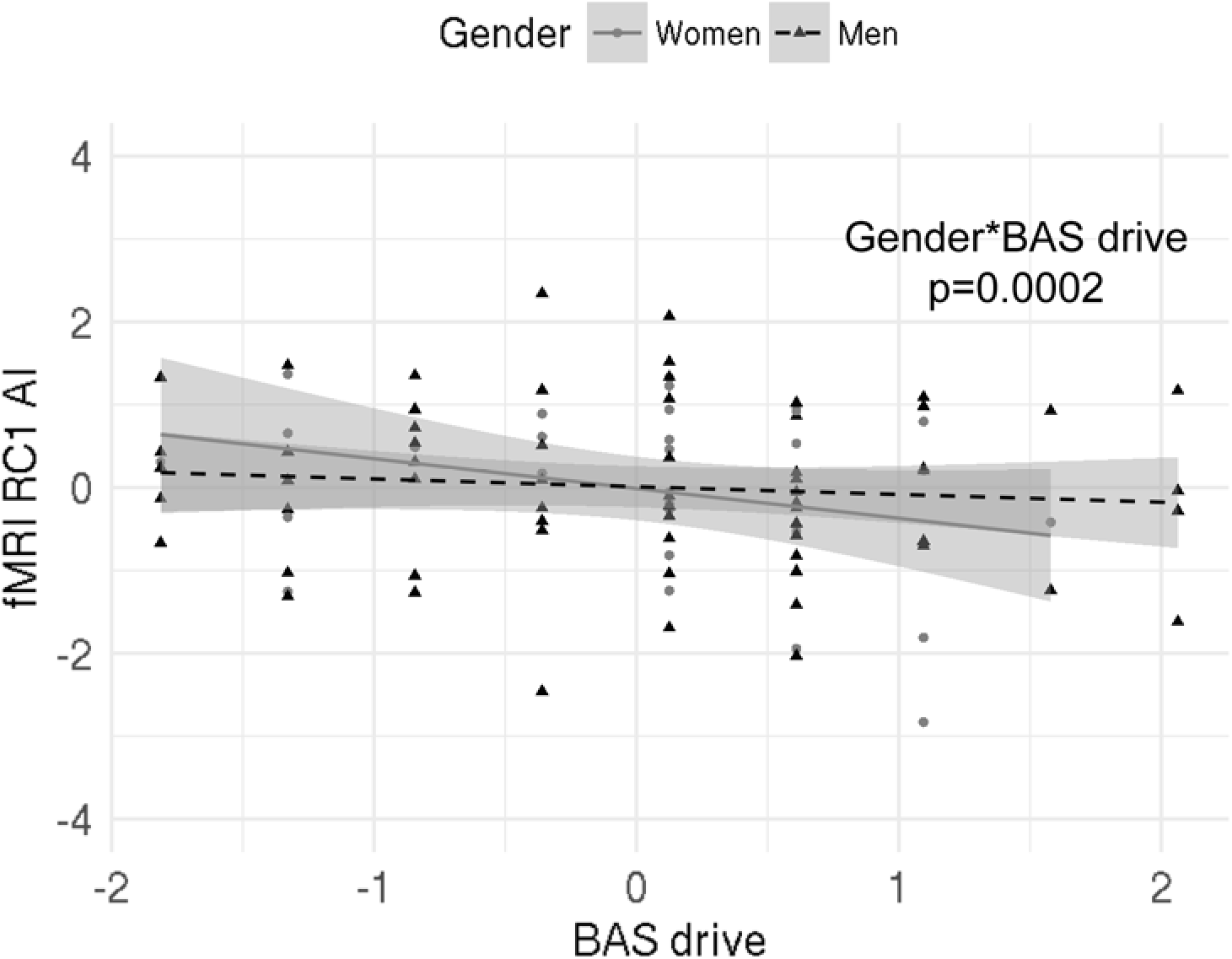
Relationship between RC1 and BAS drive scores; there was a significant interaction effect of BAS drive scores and gender on RC1, and a significant effect of BAS drive scores on RC1 in women. Index used: (L-R)/(L+R). Triangles/dots represent data points, dashed/bold lines represent the best fit, and grey shaded areas are 95% confidence intervals. RC – rotated component, AI – asymmetry index, R – right, L – left.

**Figure 3.**
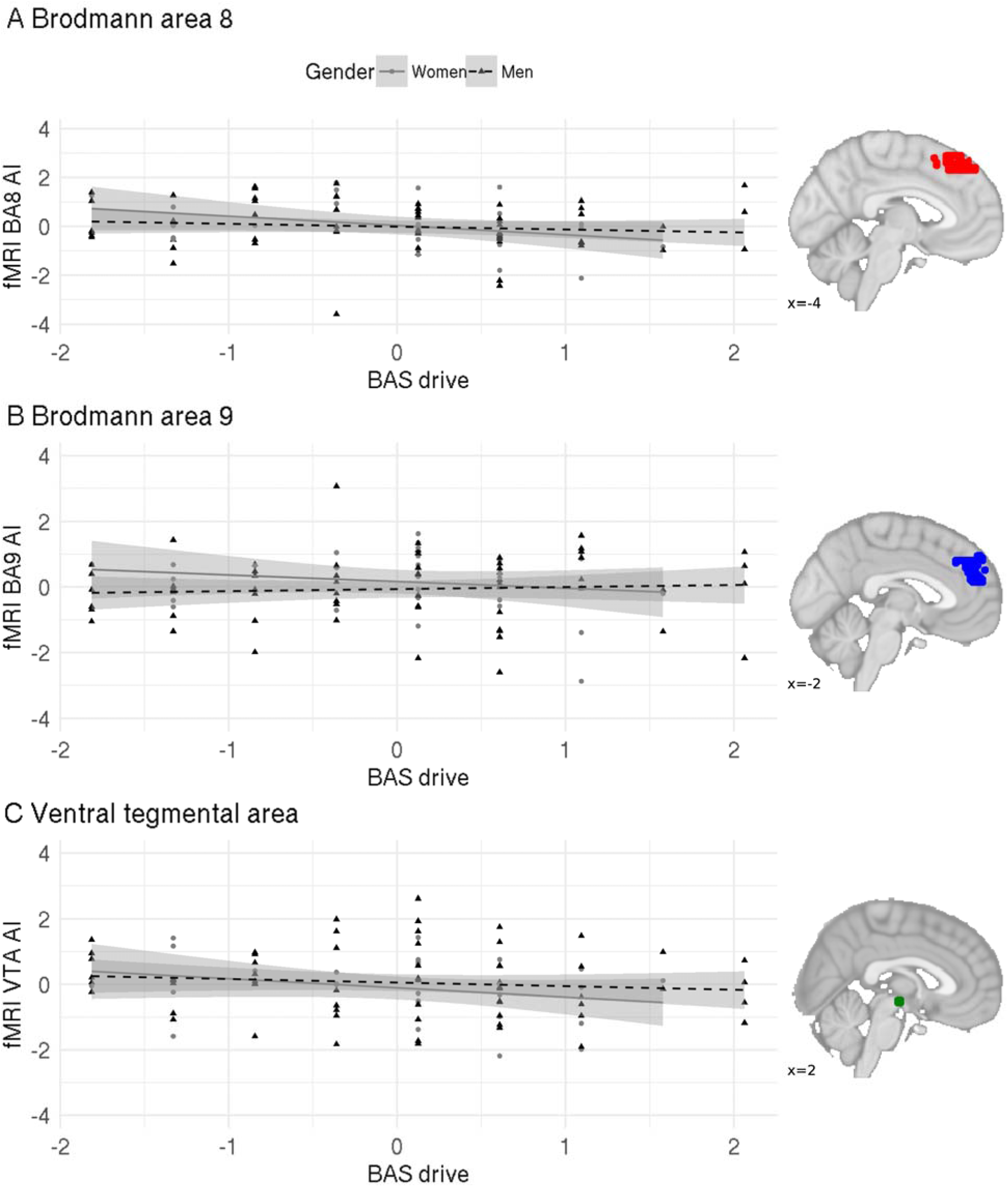
Relationship between BAS drive scores and **A** Brodmann area (BA) 8; **B** BA9; **C** ventral tegmental area (VTA). The relationship between BAS drive and a rotated component representing those three regions of interest was significant for females. Black dots represent data points, bold lines represent the best fit, and grey shaded areas are 95% confidence intervals. AI – asymmetry index, R – right, L – left.

Further, we investigated whether hemispheric asymmetries measured with fMRI are related to self-reported eating behaviour (TFEQ). This analysis included cognitive control, disinhibition and their interactions with gender as predictor variables, while the outcome variables were the 5 rotated components from the PCA analysis. Variables of no interest were BMI and age. Here, we did not find any significant relationships. Results of this analysis can be found in Table S6 and Table S7.

### 3.3 Aim 3: fMRI investigations in samples including participants with obesity - relationship of hemispheric bias and self-reported behaviours

Here, we investigated relationships between fMRI relative asymmetry indices (L-R)/(L+R) and approach/avoidance behaviours in Sample 2, characterized by a wider BMI range including individuals with overweight and obesity. The analysis included the 5 retained components describing asymmetry data as outcome variables and questionnaire variables – BAS fun, BAS drive, BAS reward responsivity, BAS – BIS, their interactions with gender, and BMI as predictors. Additionally, we included age as a regressor of no interest.

We did not find any significant relationships between approach/avoidance behaviours and fMRI hemispheric asymmetries in this sample. Results of this analysis can be found in Table S8. Loadings of each of the rotated components in the PCA can be found in Table S9.

Further, we investigated whether relative hemispheric asymmetries measured with fMRI (L-R)/(L+R) are related to self-reported eating behaviour in Samples 2 and 3 (characterised by a wider BMI range). These analyses included cognitive control, disinhibition, their interactions with gender and BMI as predictor variables, while the outcome variables were 5 rotated components from the PCA. Age was entered as a regressor of no interest. Our analyses revealed no relationships between hemispheric asymmetries and eating behaviour in both samples. Details of these analyses can be found in Tables S9-S12.

## 4 Discussion

In this study we aimed at replicating previous EEG findings concerning relationships of resting-state hemispheric asymmetries and approach/avoidance behaviours in healthy participants. Second, we aimed to investigate whether EEG asymmetry findings and fMRI asymmetry findings correspond to each other in the approach/avoidance context, as they do in the language (e.g. syntactic and semantic processing), or attention context (e.g. object or face perception) (Chakrabarty et al., 2017; Mazza & Pagano, 2017; Powell et al., 2006). Importantly, we also used fMRI to obtain data from subcortical structures, which are not easily obtainable from the EEG measures. This is an important addition especially in the context of obesity, since alterations in functions and structure of subcortical dopaminergic regions were previously often related to obesity (Cone et al., 2013; Friend et al., 2016; Geiger et al., 2009; Horstmann et al., 2015; Narayanaswami et al., 2013; Stice et al., 2011; Volkow et al., 2008; Vucetic et al., 2012). Further, we attempted to expand the findings to self-reported eating behaviour and BMI (which has been related to increased approach behaviour; Mehl et al., 2019; Mehl et al., 2018) using rsfMRI. We tested 3 independent samples: In Sample 1, we were not able to directly replicate previous EEG findings showing a positive association between BAS – BIS scores (describing individual differences between approach and avoidance behaviours) and higher left resting-state hemispheric bias. However, we show a conceptual replication of this bias with BAS drive in women. Second, we show that BAS drive scores are related to asymmetries measured by rsfMRI – with an opposite relationship to the one found in EEG. Further, in Sample 2 – that included participants with overweight and obesity as well as rsfMRI data – we did not find any relationship of hemispheric bias and approach/avoidance behaviour or BMI using the same measures as in Sample 1. Finally, in none of the samples did we find relationships of hemispheric bias and self-reported eating behaviour.

Past work by Gray and colleagues has suggested that human behaviour is driven by the interplay of the behavioural inhibition and activation systems (Gray, 1981; Gray & McNaughton, 1992). In a number of clinical and laboratory studies, it has been proposed that those fundamental behavioural dimensions are driven by asymmetric engagements of anterior brain regions (Davidson & Hugdahl, 1995). In particular, the neural substrate for the inhibition system or withdrawal behaviour was found in the right prefrontal cortex, while the left prefrontal cortex was related to approach behaviour (Davidson & Hugdahl, 1995). Those conclusions are based predominantly on rsEEG studies but also on studies in patients with frontal brain lesions. In our work we aimed to replicate the seminal study by Sutton and Davidson (1997), which showed a positive association of BAS – BIS differential scores with left hemispheric bias, as measured by absolute alpha power from rsEEG. Although we have analysed our data in the same way, we did not replicate these results. In our study, the rsEEG duration was 16 minutes (eyes closed + eyes open) as opposed to 8 minutes in Sutton’s study (eyes closed + eyes open), however, longer duration might provide a better estimation of resting-state processes. Yet it is unlikely that those small methodological differences can explain lack of direct replication. However, our sample size was much larger and included participants in a wider age-range. Additionally, gender distribution was not equal, whereas in Sutton’s study 50% of the sample were women (although we statistically controlled for age, BMI and gender). Those factors might influence results beyond what is possible to be corrected by means of statistical analysis (as we statistically controlled for age, BMI, and gender).

Importantly, in a more detailed EEG data analysis using refined a relative asymmetry index, that is superior to an absolute in terms of interpretability, and relative alpha power, we found effects that are conceptually similar to the ones by Sutton and Davidson (1997): we found a positive gender-specific relationship between left hemispheric bias (indicating increased left over right hemispheric activity) and BAS drive. Additional analyses showed that this effect is specific for the eyes open condition, as an identical analysis with the mean of EO and EC conditions as outcome variables did not show any significant results. It is conceivable that BIS/BAS correlates only with EEG hemispheric asymmetries during an EO resting condition because approach/avoidance behaviours inherently require engaging with the environment in order to perceive and react to stimuli.

The effect observed in EEG analysis in eyes open condition indicates that higher approach behaviour (or drive towards positive reinforcement) is related to higher left brain activity at rest. While Sutton and Davidson (1997) found a similar association in a sample including both genders, in our sample it was only true for women. As Sutton and Davidson did not explicitly test gender differences, it cannot be excluded that their findings were driven by women. Further, in this study we found significant effects using a different measure of approach behaviour (BAS drive versus BAS – BIS scores). BAS drive describes an absolute strength of the approach system (drive towards positive stimuli), while BAS – BIS difference scores represent the balance between the two systems. It is possible that those different measures are related to hemispheric asymmetries in a distinct, gender-dependent way. Nevertheless, previous literature shows that gender indeed influences hemispheric asymmetries – brains of men seem to be more lateralised as compared to women (Hausmann, 2002, 2017; McGlone, 1980). This does not exclude the possibility that women’s brains show different associations between hemispheric asymmetries and self-reported behaviours, possibly through sex hormones (Hausmann, 2002, 2017). Future studies should aim to replicate our result and investigate asymmetries specifically with regard to gender differences.

It is worth noting that we found significant associations of questionnaire measures and hemispheric asymmetries measured with low relative alpha power, but not with broadband relative alpha power. Since low alpha power represents such attentional processes as vigilance (Klimesch et al., 2007; Petsche et al., 1997), our results suggest that hemispheric asymmetries are related to those processes, rather than to general inhibitory processing within the brain. In the EEG analysis in Sample 1, we also found a significant effect of BMI, with increased BMI being related to higher left vs. right hemispheric activity. However, Sample 1 included predominantly lean participants and the findings cannot be interpreted in relation to individuals with overweight or obesity, but rather to the variance in BMI within the normal range.

The second aim of our study was to investigate whether approach/avoidance-related asymmetries can be measured with both EEG and fMRI. We show that the relationship between hemispheric asymmetries, as measured by fMRI and fALFF, and BAS drive, is opposite to the one found in the EEG data. This is interesting for two reasons: First, it suggests that there might be an indirect relationship between two fundamentally different (Scheeringa et al., 2011) measures of brain activity (by means of correlations with the same behavioural measures). This is even despite the fact that we did not find a direct correlation between the two measures. Second, it provides evidence that fMRI measures of hemispheric asymmetry can be related to approach and avoidance behaviours. This provides additional methodological possibilities to investigate relationships between hemispheric asymmetries and behavioural measures of approach/avoidance. Interestingly, the direction of the relationships measured by EEG and fMRI were opposite. This indicates that alpha power and fALFF might measure different processes, which is also reflected in a lack of direct relationship between EEG and whole-brain fALFF asymmetries. Alpha power indeed is conceptualised to be inversely related to brain activity by enabling active inhibition (Klimesch et al., 2007). fALFF, on the other hand, is generally suggested to be a measure of brain activity (Zou et al., 2008).

We therefore hypothesised that alpha power and fALFF could simply be inversely related to each other. This is not supported by our data. Instead, this relationship seems to be more complex. In our correlation of relative alpha power and fALFF we did not find any evidence for such a relationship. This might be because EEG and fMRI measure electrical activity and hemodynamic response, respectively, but also because the oscillations measured by the two methods differ greatly in frequency ranges (8-12Hz vs. 0.01-0.1Hz). It is nevertheless encouraging that the asymmetries measured with fMRI and EEG show relationships to the same behavioural measures. This provides a first step for further investigations of relationships between relative alpha power and fALFF measurements, ideally using combined EEG-fMRI.

Finding relationships between fMRI asymmetry measures and behaviour, we focused on the third aim of the study – investigating this relationship in fMRI-only samples including participants with overweight and obesity, where unfortunately only fMRI data were available. Concerning approach and avoidance behaviours, we used data of a sample which included lean, overweight and obese people. We investigated relationships between hemispheric bias and BIS/BAS questionnaires. Additionally, we investigated a direct relationship between hemispheric bias and BMI, since BMI in the obese range is related to increased approach behaviour (Mehl et al., 2018), and obesity has been described as a deficiency of right-brain activation (Alonso-Alonso & Pascual-Leone, 2007). Our analyses did not show a significant relationship between hemispheric bias and BMI or between hemispheric bias and approach/avoidance behaviour. Thus, we did not find support for the right-brain theory of obesity, which suggests that hemispheric biases at rest may not be related to BMI *per se*, but to specific patterns of approach/avoidance and/or eating behaviour instead. Relatedly, it is conceivable that hemispheric biases during specific task performance might be related to BMI. While previous studies supporting the right brain theory of obesity largely focused on patients with unilateral brain lesions or structural asymmetries (Colcombe et al., 2006; Regard & Landis, 1997; Short et al., 2005; Uher & Treasure, 2005), our resting-state data were obtained in neurologically healthy participants. This may imply that previous results on obesity-related hemispheric asymmetries cannot be generalized to individuals with obesity. This heterogeneity, while increasing ecological validity, might introduce noise, which in turn makes it difficult to detect associations between BMI and hemispheric asymmetries. Lastly, the right-brain theory of obesity is based on a number of findings relating eating behaviours and physical activity to hemispheric asymmetries, and not to BMI directly (Colcombe et al., 2006; Regard & Landis, 1997; Short et al., 2005; Uher & Treasure, 2005), as did our study – which might explain deviating results. In sum, future studies need to focus on relationships between obesity measures and hemispheric asymmetries in EEG and fMRI measurements of both resting-state and task contexts to confirm or revise the right-brain theory of obesity.

We further investigated associations between hemispheric asymmetries and self-reported eating behaviours in all 3 samples. Here, we did not find any relationships using rsEEG and rsfMRI data. That is, we were not able to replicate previous rsEEG findings showing hemispheric bias relationships with disinhibition, hunger (Ochner et al., 2009), or restrained eating (Silva et al., 2002). Similarly, the study by Ochner and colleagues (2009) included participants with overweight and obesity (so did 2 of our 3 samples), and the study by Silva and colleagues (2002) included only lean women (one of our samples included mostly lean participants and we investigated interactions with gender). However, certain differences between those studies and our research exist, which might explain different results: First, Ochner and colleagues investigated a group of much older participants (mean age: 49 years). It is conceivable that the duration of obesity influences prefrontal asymmetries, hence age might explain differences between results. Furthermore, in our study we were very conservative with regard to multiple comparisons correction, while Ochner and colleagues were more liberal in this respect.

Overall, the measure of hemispheric asymmetries utilising fALFF and the relationship of this measure with approach/avoidance behaviour seem to be unstable and highly dependent on the characteristics of samples under study, predominantly the BMI distribution. Further, the relationship between hemispheric asymmetries and approach/avoidance behaviour seems to be dependent on the way in which this behaviour is measured, that is, the questionnaire used. More research is needed to investigate which different behavioural measures influence this relationship. One way to improve current research is to use large and well-characterised publicly available datasets.

Some limitations of our study need to be acknowledged: EEG data were only available for one sample. It would provide additional evidence to investigate differences between rsEEG and rsfMRI asymmetry associations with behavioural measures in other samples, especially concerning BMI and eating behaviour – aspects not investigated as thoroughly as approach/avoidance behaviours. As our study investigated relationships between self-reported approach/avoidance behaviours and resting-state neuroimaging measures, future studies could also include task-based neuroimaging measures, especially in the context of obesity. This might give a more valid proxy for everyday motivational behaviours and therefore have higher ecological validity.

In sum, we conceptually replicated findings showing relationships between hemispheric bias and approach/avoidance behaviours in women, but not self-reported eating behaviour in both rsEEG and rsfMRI. Moreover, we investigated relationships between rsEEG alpha power measures and rsfMRI fALFF. We show that associations of hemispheric asymmetries measured with rsEEG and rsfMRI in terms of approach/avoidance behaviours are opposite. Future studies should answer the question of how those measures relate to each other in a more systematic way. We suggest that future studies should be performed using samples of lean, overweight and obese participants using both EEG and fMRI measures.

## Acknowledgments

This work was supported by the German Federal Ministry of Education and Research (BMBF) to MG (FKZ: 13GWl0206B) and in the framework of the Integrated Research and Treatment Center AdiposityDiseases at the University of Leipzig (FKZ: 01EO1001) to FM, LJ, AH, JN; the University Research Priority Program “Dynamics of Healthy Aging” of the University of Zurich to FL; German Research Foundation, Collaborative Research Centre 1052 ‘Obesity Mechanisms’, subproject A5, at the University of Leipzig to AH.

## References

Aberg, K. C., Doell, K. C., & Schwartz, S. (2015). Hemispheric Asymmetries in Striatal Reward Responses Relate to Approach–Avoidance Learning and Encoding of Positive–Negative Prediction Errors in Dopaminergic Midbrain Regions. The Journal of Neuroscience, 35(43), 14491–14500. doi:10.1523/JNEUROSCI.1859-15.2015

Adcock, R. A., Thangavel, A., Whitfield-Gabrieli, S., Knutson, B., & Gabrieli, J. D. E. (2006). Reward-Motivated Learning: Mesolimbic Activation Precedes Memory Formation. Neuron, 50(3), 507–517. doi:10.1016/j.neuron.2006.03.036

Alonso-Alonso, M., & Pascual-Leone, A. (2007). The Right Brain Hypothesis for Obesity. JAMA, 297(16), 1819–1822. doi:10.1001/jama.297.16.1819

Avants, B. B., Tustison, N. J., Song, G., Cook, P. A., Klein, A., & Gee, J. C. (2011). A reproducible evaluation of ANTs similarity metric performance in brain image registration. NeuroImage, 54(3), 2033–2044. doi:10.1016/j.neuroimage.2010.09.025

Babayan, A., Erbey, M., Kumral, D., Reinelt, J. D., Reiter, A. M. F., Röbbig, J., Villringer, A. (2019). A mind-brain-body dataset of MRI, EEG, cognition, emotion, and peripheral physiology in young and old adults. Scientific Data, 6, 180308. doi:10.1038/sdata.2018.308

Babiloni, C., Marzano, N., Lizio, R., Valenzano, A., Triggiani, A. I., Petito, A., Del Percio, C. (2011). Resting state cortical electroencephalographic rhythms in subjects with normal and abnormal body weight. NeuroImage, 58(2), 698–707. doi:10.1016/j.neuroimage.2011.05.080

Bazanova, O. M. (2012). Alpha EEG Activity Depends on the Individual Dominant Rhythm Frequency. Journal of Neurotherapy, 16(4), 270–284. doi:10.1080/10874208.2012.730786

Bazanova, O. M., & Vernon, D. (2014). Interpreting EEG alpha activity. Neuroscience and Biobehavioral Reviews, 44, 94–110. doi:10.1016/j.neubiorev.2013.05.007

Beck, A. T., Steer, R. A., Ball, R., & Ranieri, W. F. (1996). Comparison of Beck Depression Inventories-IA and-II in Psychiatric Outpatients. Journal of Personality Assessment, 67(3), 588–597. doi:10.1207/s15327752jpa6703_13

Behzadi, Y., Restom, K., Liau, J., & Liu, T. T. (2007). A Component Based Noise Correction Method (CompCor) for BOLD and Perfusion Based fMRI. NeuroImage, 37(1), 90–101. doi:10.1016/j.neuroimage.2007.04.042

Bell, A. J., & Sejnowski, T. J. (1995). An Information-Maximization Approach to Blind Separation and Blind Deconvolution. Neural Computation, 7(6), 1129–1159. doi:10.1162/neco.1995.7.6.1129

Biswal, B., Yetkin, F. Z., Haughton, V. M., & Hyde, J. S. (1995). Functional connectivity in the motor cortex of resting human brain using echo-planar mri. Magnetic Resonance in Medicine, 34(4), 537–541. doi:10.1002/mrm.1910340409

Carver, C. S., & White, T. L. (1994). Behavioral inhibition, behavioral activation, and affective responses to impending reward and punishment: The BIS/BAS Scales. Journal of Personality and Social Psychology, 67(2), 319–333. doi:10.1037/0022-3514.67.2.319

Chakrabarty, M., Badgio, D., Ptacek, J., Biswas, A., Ghosal, M., & Chatterjee, G. (2017, 2017). Hemispheric Asymmetry in Attention and its Impact on Our Consciousness: A Review with Reference to Altered Consciousness in Right Hemisphere Damaged Subjects.

Colcombe, S. J., Erickson, K. I., Scalf, P. E., Kim, J. S., Prakash, R., McAuley, E., Kramer, A. F. (2006). Aerobic exercise training increases brain volume in aging humans. The Journals of Gerontology. Series A, Biological Sciences and Medical Sciences, 61(11), 1166–1170.

Cone, J. J., Chartoff, E. H., Potter, D. N., Ebner, S. R., & Roitman, M. F. (2013). Prolonged High Fat Diet Reduces Dopamine Reuptake without Altering DAT Gene Expression. PLOS ONE, 8(3), e58251. doi:10.1371/journal.pone.0058251

Cousijn, J., Goudriaan, A. E., Ridderinkhof, K. R., Brink, W. v. d., Veltman, D. J., & Wiers, R. W. (2012). Approach-Bias Predicts Development of Cannabis Problem Severity in Heavy Cannabis Users: Results from a Prospective FMRI Study. PLOS ONE, 7(9), e42394. doi:10.1371/journal.pone.0042394

Davidson, R. J. (1993). Cerebral asymmetry and emotion: Conceptual and methodological conundrums. Cognition and Emotion, 7(1), 115–138. doi:10.1080/02699939308409180

Davidson, R. J. (1994). Asymmetric brain function, affective style, and psychopathology: The role of early experience and plasticity. Development and Psychopathology, 6(04), 741–758. doi:10.1017/S0954579400004764

Davidson, R. J., & Hugdahl, K. (1995). Brain Asymmetry. Cambridge, Massachusetts: The MIT Press.

Delorme, A., & Makeig, S. (2004). EEGLAB: an open source toolbox for analysis of single-trial EEG dynamics including independent component analysis. Journal of Neuroscience Methods, 134(1), 9–21. doi:10.1016/j.jneumeth.2003.10.009

Dietrich, A., Federbusch, M., Grellmann, C., Villringer, A., & Horstmann, A. (2014). Body weight status, eating behavior, sensitivity to reward/punishment, and gender: relationships and interdependencies. Eating Behavior, 5, 1073. doi:10.3389/fpsyg.2014.01073

Fransson, P. (2005). Spontaneous low-frequency BOLD signal fluctuations: an fMRI investigation of the resting-state default mode of brain function hypothesis. Human Brain Mapping, 26(1), 15–29. doi:10.1002/hbm.20113

Friend, D. M., Devarakonda, K., O’Neal, T. J., Skirzewski, M., Papazoglou, I., Kaplan, A. R., Kravitz, A. V. (2016). Basal Ganglia Dysfunction Contributes to Physical Inactivity in Obesity. Cell Metabolism, 0(0). doi:10.1016/j.cmet.2016.12.001

Geiger, B. M., Haburcak, M., Avena, N. M., Moyer, M. C., Hoebel, B. G., & Pothos, E. N. (2009). Deficits of mesolimbic dopamine neurotransmission in rat dietary obesity. Neuroscience, 159(4), 1193–1199. doi:10.1016/j.neuroscience.2009.02.007

Giacometti, P., Perdue, K. L., & Diamond, S. G. (2014). Algorithm to find high density EEG scalp coordinates and analysis of their correspondence to structural and functional regions of the brain. Journal of Neuroscience Methods, 229, 84–96. doi:10.1016/j.jneumeth.2014.04.020

Gorgolewski, K., Burns, C. D., Madison, C., Clark, D., Halchenko, Y. O., Waskom, M. L., & Ghosh, S. S. (2011). Nipype: A Flexible, Lightweight and Extensible Neuroimaging Data Processing Framework in Python. Frontiers in Neuroinformatics, 5. doi:10.3389/fninf.2011.00013

Gray, J. A. (1981). A Critique of Eysenck’s Theory of Personality. In A Model for Personality (pp. 246–276): Springer, Berlin, Heidelberg.

Gray, J. A., & McNaughton, N. (1992). The Neuropsychology of Anxiety: An Enquiry into the Functions of the Septo-Hippocampal System (Oxford Psychology Series): An Enquiry into the Function of the Septo-hippocampal System (Second ed.). Oxford New York: Oxford University Press, USA.

Greve, D. N., & Fischl, B. (2009). Accurate and robust brain image alignment using boundary-based registration. NeuroImage, 48(1), 63–72. doi:10.1016/j.neuroimage.2009.06.060

Hausmann, M. (2002). Functional cerebral asymmetries during the menstrual cycle: a cross-sectional and longitudinal analysis. 40(7), 808–816. doi:10.1016/s0028-3932(01)00179-8

Hausmann, M. (2017). Why sex hormones matter for neuroscience: A very short review on sex, sex hormones, and functional brain asymmetries. 95(1-2), 40–49. doi:10.1002/jnr.23857

Herwig, U., Satrapi, P., & Schönfeldt-Lecuona, C. (2003). Using the International 10-20 EEG System for Positioning of Transcranial Magnetic Stimulation. Brain Topography, 16(2), 95–99. doi:10.1023/B:BRAT.0000006333.93597.9d

Hiroshige, Y., & Dorokhov, V. B. (1997). Hemispheric Asymmetry and Regional Differences in Electroencephalographic Alpha Activity at the Wake-Sleep Transition. Japanese Psychological Research, 39(2), 75–86. doi:10.1111/1468-5884.00041

Hoaglin, D. C., & Iglewicz, B. (1987). Fine-Tuning Some Resistant Rules for Outlier Labeling. Journal of the American Statistical Association, 82(400), 1147–1149. doi:10.1080/01621459.1987.10478551

Hoaglin, D. C., Iglewicz, B., & Tukey, J. W. (1986). Performance of Some Resistant Rules for Outlier Labeling. Journal of the American Statistical Association, 81(396), 991–999. doi:10.1080/01621459.1986.10478363

Horstmann, A., Fenske, W. K., & Hankir, M. K. (2015). Argument for a non-linear relationship between severity of human obesity and dopaminergic tone. Obesity Reviews, n/a-n/a. doi:10.1111/obr.12303

Jenkinson, M., Bannister, P., Brady, M., & Smith, S. (2002). Improved Optimization for the Robust and Accurate Linear Registration and Motion Correction of Brain Images. NeuroImage, 17(2), 825–841. doi:10.1006/nimg.2002.1132

Jenkinson, M., Beckmann, C. F., Behrens, T. E. J., Woolrich, M. W., & Smith, S. M. (2012). FSL. NeuroImage, 62(2), 782–790. doi:10.1016/j.neuroimage.2011.09.015

Johnson, S. L., Turner, R. J., & Iwata, N. (2003). BIS/BAS levels and psychiatric disorder: An epidemiological study. Journal of Psychopathology and Behavioral Assessment, 25(1), 25–36. doi:10.1023/A:1022247919288

Jolliffe, I. T. (2002). Principal Component Analysis (2 ed.). New York: Springer-Verlag.

Jolliffe, I. T., & Cadima, J. (2016). Principal component analysis: a review and recent developments. Philosophical transactions. Series A, Mathematical, physical, and engineering sciences, 374(2065). doi:10.1098/rsta.2015.0202

Kiviniemi, V., Jauhiainen, J., Tervonen, O., Pääkkö, E., Oikarinen, J., Vainionpää, V., Biswal, B. (2000). Slow vasomotor fluctuation in fMRI of anesthetized child brain. Magnetic Resonance in Medicine, 44(3), 373–378.

Klimesch, W., Sauseng, P., & Hanslmayr, S. (2007). EEG alpha oscillations: The inhibition–timing hypothesis. Brain Research Reviews, 53(1), 63–88. doi:10.1016/j.brainresrev.2006.06.003

Liem, F., Varoquaux, G., Kynast, J., Beyer, F., Kharabian Masouleh, S., Huntenburg, J. M., Margulies, D. S. (2017). Predicting brain-age from multimodal imaging data captures cognitive impairment. NeuroImage, 148, 179–188. doi:10.1016/j.neuroimage.2016.11.005

Maldjian, J. A., Laurienti, P. J., Kraft, R. A., & Burdette, J. H. (2003). An automated method for neuroanatomic and cytoarchitectonic atlas-based interrogation of fMRI data sets. NeuroImage, 19(3), 1233–1239. doi:10.1016/S1053-8119(03)00169-1

Mathar, D., Wilkinson, L., Holl, A. K., Neumann, J., Deserno, L., Villringer, A., Horstmann, A. (2017). The role of dopamine in positive and negative prediction error utilization during incidental learning – Insights from Positron Emission Tomography, Parkinson’s disease and Huntington’s disease. Cortex, 90, 149–162. doi:10.1016/j.cortex.2016.09.004

Mazza, V., & Pagano, S. (2017). Electroencephalographic asymmetries in human cognition. Neuromethods, 122, 407–439. doi:10.1007/978-1-4939-6725-4_13

McGlone, J. (1980). Sex differences in human brain asymmetry: a critical survey. Behavioral and Brain Sciences, 3(2), 215–227. doi:10.1017/S0140525X00004398

Mehl, N., Morys, F., Villringer, A., & Horstmann, A. (2019). Unhealthy yet Avoidable—How Cognitive Bias Modification Alters Behavioral and Brain Responses to Food Cues in Individuals with Obesity. Nutrients, 11(4), 874. doi:10.3390/nu11040874

Mehl, N., Mueller-Wieland, L., Mathar, D., & Horstmann, A. (2018). Retraining automatic action tendencies in obesity. Physiology & Behavior. doi:10.1016/j.physbeh.2018.03.031

Mendes, N., Oligschlaeger, S., Lauckner, M. E., Golchert, J., Huntenburg, J. M., Falkiewicz, M., … Margulies, D. S. (2017). A functional connectome phenotyping dataset including cognitive state and personality measures. bioRxiv, 164764. doi:10.1101/164764

Morgan, B. E., Van, H., Hermans, E. J., Scholten, M. R. M., Stein, D. J., & Kahn, R. S. (2009). Gray’s BIS/BAS dimensions in non-comorbid, non-medicated social anxiety disorder. World Journal of Biological Psychiatry, 10(4 PART 3), 925–928. doi:10.1080/15622970802571695

Narayanaswami, V., Thompson, A. C., Cassis, L. A., Bardo, M. T., & Dwoskin, L. P. (2013). Diet-induced obesity: dopamine transporter function, impulsivity and motivation. International Journal of Obesity, 37(8), 1095–1103. doi:10.1038/ijo.2012.178

Neto, L. L., Oliveira, E., Correia, F., & Ferreira, A. G. (2008). The human nucleus accumbens: where is it? A stereotactic, anatomical and magnetic resonance imaging study. Neuromodulation: Journal of the International Neuromodulation Society, 11(1), 13–22. doi:10.1111/j.1525-1403.2007.00138.x

Ochner, C. N., Green, D., van Steenburgh, J. J., Kounios, J., & Lowe, M. R. (2009). Asymmetric prefrontal cortex activation in relation to markers of overeating in obese humans. Appetite, 53(1), 44–49. doi:10.1016/j.appet.2009.04.220

Oostenveld, R., & Praamstra, P. (2001). The five percent electrode system for high-resolution EEG and ERP measurements. Clinical Neurophysiology: Official Journal of the International Federation of Clinical Neurophysiology, 112(4), 713–719.

Petsche, H., Kaplan, S., von Stein, A., & Filz, O. (1997). The possible meaning of the upper and lower alpha frequency ranges for cognitive and creative tasks. International Journal of Psychophysiology: Official Journal of the International Organization of Psychophysiology, 26(1-3), 77–97.

Pivik, R. T., Broughton, R. J., Coppola, R., Davidson, R. J., Fox, N., & Nuwer, M. R. (1993). Guidelines for the recording and quantitative analysis of electroencephalographic activity in research contexts. Psychophysiology, 30(6), 547–558.

Powell, H. W. R., Parker, G. J. M., Alexander, D. C., Symms, M. R., Boulby, P. A., Wheeler-Kingshott, C. A. M., Duncan, J. S. (2006). Hemispheric asymmetries in language-related pathways: A combined functional MRI and tractography study. NeuroImage, 32(1), 388–399. doi:10.1016/j.neuroimage.2006.03.011

Power, J. D., Barnes, K. A., Snyder, A. Z., Schlaggar, B. L., & Petersen, S. E. (2012). Spurious but systematic correlations in functional connectivity MRI networks arise from subject motion. NeuroImage, 59(3), 2142–2154. doi:10.1016/j.neuroimage.2011.10.018

Regard, M., & Landis, T. (1997). “Gourmand syndrome”: eating passion associated with right anterior lesions. Neurology, 48(5), 1185–1190.

Richman, M. B. (1986). Rotation of principal components. Journal of Climatology, 6(3), 293–335. doi:10.1002/joc.3370060305

Richman, M. L. B. (1987). Rotation of principal components: A reply. Journal of Climatology, 7(5), 511–520. doi:10.1002/joc.3370070507

Scheeringa, R., Fries, P., Petersson, K.-M., Oostenveld, R., Grothe, I., Norris, D. G., Bastiaansen, M. C. M. (2011). Neuronal dynamics underlying high- and low-frequency EEG oscillations contribute independently to the human BOLD signal. Neuron, 69(3), 572–583. doi:10.1016/j.neuron.2010.11.044

Short, R. A., Broderick, D. F., Patton, A., Arvanitakis, Z., & Graff-Radford, N. R. (2005). Different patterns of magnetic resonance imaging atrophy for frontotemporal lobar degeneration syndromes. Archives of Neurology, 62(7), 1106–1110. doi:10.1001/archneur.62.7.1106

Silva, J. R., Pizzagalli, D. A., Larson, C. L., Jackson, D. C., & Davidson, R. J. (2002). Frontal brain asymmetry in restrained eaters. Journal of Abnormal Psychology, 111(4), 676–681. doi:10.1037/0021-843X.111.4.676

Stice, E., Yokum, S., Burger, K. S., Epstein, L. H., & Small, D. M. (2011). Youth at Risk for Obesity Show Greater Activation of Striatal and Somatosensory Regions to Food. The Journal of Neuroscience, 31(12), 4360–4366. doi:10.1523/JNEUROSCI.6604-10.2011

Stunkard, A. J., & Messick, S. (1985). The three-factor eating questionnaire to measure dietary restraint, disinhibition and hunger. Journal of Psychosomatic Research, 29(1), 71–83. doi:10.1016/0022-3999(85)90010-8

Sutton, S. K., & Davidson, R. J. (1997). Prefrontal Brain Asymmetry: A Biological Substrate of the Behavioral Approach and Inhibition Systems. Psychological Science, 8(3), 204–210. doi:10.1111/j.1467-9280.1997.tb00413.x

Tomer, R., Goldstein, R. Z., Wang, G.-J., Wong, C., & Volkow, N. D. (2008). Incentive motivation is associated with striatal dopamine asymmetry. Biological Psychology, 77(1), 98–101. doi:10.1016/j.biopsycho.2007.08.001

Tomer, R., Slagter, H. A., Christian, B. T., Fox, A. S., King, C. R., Murali, D., Davidson, R. J. (2013). Love to Win or Hate to Lose? Asymmetry of Dopamine D2 Receptor Binding Predicts Sensitivity to Reward versus Punishment. Journal of Cognitive Neuroscience, 26(5), 1039–1048. doi:10.1162/jocn_a_00544

Towle, V. L., Bolaños, J., Suarez, D., Tan, K., Grzeszczuk, R., Levin, D. N., Spire, J.-P. (1993). The spatial location of EEG electrodes: locating the best-fitting sphere relative to cortical anatomy. Electroencephalography and Clinical Neurophysiology, 86(1), 1–6. doi:10.1016/0013-4694(93)90061-Y

Tukey, J. W. (1977). Exploratory Data Analysis (1 edition ed.). Reading, Mass: Pearson.

Uher, R., & Treasure, J. (2005). Brain lesions and eating disorders. Journal of Neurology, Neurosurgery, and Psychiatry, 76(6), 852–857. doi:10.1136/jnnp.2004.048819

Volkow, N. D., Wang, G.-J., Fowler, J. S., & Telang, F. (2008). Overlapping Neuronal Circuits in Addiction and Obesity: Evidence of Systems Pathology. Philosophical Transactions: Biological Sciences, 363(1507), 3191–3200.

Vucetic, Z., Carlin, J. L., Totoki, K., & Reyes, T. M. (2012). Epigenetic dysregulation of the dopamine system in diet-induced obesity. Journal of Neurochemistry, 120(6), 891–898. doi:10.1111/j.1471-4159.2012.07649.x

Wiers, C. E., Kühn, S., Javadi, A. H., Korucuoglu, O., Wiers, R. W., Walter, H., Bermpohl, F. (2013). Automatic approach bias towards smoking cues is present in smokers but not in ex-smokers. Psychopharmacology, 229(1), 187–197. doi:10.1007/s00213-013-3098-5

Wiers, C. E., Stelzel, C., Park, S. Q., Gawron, C. K., Ludwig, V. U., Gutwinski, S., Bermpohl, F. (2014). Neural Correlates of Alcohol-Approach Bias in Alcohol Addiction: the Spirit is Willing but the Flesh is Weak for Spirits. Neuropsychopharmacology, 39(3), 688–697. doi:10.1038/npp.2013.252

Zou, Q.-H., Zhu, C.-Z., Yang, Y., Zuo, X.-N., Long, X.-Y., Cao, Q.-J., Zang, Y.-F. (2008). An improved approach to detection of amplitude of low-frequency fluctuation (ALFF) for resting-state fMRI: Fractional ALFF. Journal of Neuroscience Methods, 172(1), 137–141. doi:10.1016/j.jneumeth.2008.04.012

