## Supplementary material for "Hemispheric asymmetries in resting-state EEG and fMRI are related to approach and avoidance behaviour, but not to eating behaviour or BMI": Supplemetary materials

**Title**

**Affiliations**

### 1. Supplementary results

#### 1.1. Samples comparison

To compare the three samples regarding their demographic characteristics we ran separate one-way ANOVAs for BMI and age, and a χ^2^ test for equality of gender distribution between the samples. We followed up the ANOVAs with *post hoc* Tukey’s tests to determine which groups differed from each other. The results of these analyses can be found in Table S1. We found that regarding BMI all samples differed from each other, while regarding age, Sample 2 differed from both other samples, which did not significantly differ from each other. Concerning gender distribution, all samples differed from each other.

#### 1.2. Questionnaire data – samples comparison

Table S1 represents questionnaire data of all 3 samples included in the study. Performed ANOVAs indicated group differences in both cognitive control and disinhibition scales. Tukey’s tests showed that concerning cognitive control there were no significant pairwise differences between groups. Regarding disinhibition, Sample 2 differed significantly from both Samples 1 and 3. We further performed four ANOVAs to compare BIS/BAS data in Samples 1 and 2. This analysis revealed no significant differences for this questionnaire.

### 2. Supplementary Tables

Table S1 Means, standard deviation and statistical tests concerning questionnaire differences in all experimental samples. BIS/BAS questionnaire data were available for Samples 1 and 2, whereas TFEQ data were available for all samples. CC – cognitive control, DI – disinhibition, BAS – behavioural activation system, BIS – behavioural inhibition system.

|  | Sample 1 (n=117) | | | Sample 2 (n=89) | | | Sample 3 (n=152) | | | ANOVA | | Sample 1 vs. Sample 2 | Sample 1 vs. Sample 3 | Sample 2 vs. Sample 3 |
| --- | --- | --- | --- | --- | --- | --- | --- | --- | --- | --- | --- | --- | --- | --- |
|  | Mean | SD | Range | Mean | SD | Range | Mean | SD | Range | F(2,355)-value | p-value |  |  |  |
| BMI (kg/m^2^) | 23.01 | 2.57 | 17.95-31.80 | 29.54 | 8.25 | 17.67-59.78 | 26.40 | 5.62 | 16.26-49.96 | 33.70 | **<0.0001** | **<0.0001** | **<0.0001** | **<0.0001** |
| Age (years) | 25 | 3 | 20-35 | 27 | 4 | 20-37 | 24 | 4.5 | 18-35 | 18.08 | **<0.0001** | **<0.0001** | **<0.0001** | **<0.0001** |
| CC | 5.56 | 4.53 | 0-18 | 6.92 | 4.60 | 0-20 | 6.78 | 4.57 | 0-20 | 3.073 | **0.0480** | 0.0885 | 0.0777 | 0.9719 |
| DI | 5.23 | 2.48 | 1-12 | 6.79 | 3.52 | 1-15 | 4.63 | 2.90 | 1-15 | 15.36 | **<0.0001** | **0.0006** | 0.2158 | **<0.0001** |
|  | | | | | | | | | | F(1,204) | p-value |  | | |
| BAS fun | 3.13 | 0.41 | 2.25-4.00 | 3.03 | 0.50 | 1.75-4.00 | - | - |  | 2.213 | 0.1380 | - | - | - |
| BAS drive | 2.94 | 0.51 | 2.00-4.00 | 2.92 | 0.51 | 1.50-4.00 | - | - |  | 0.103 | 0.7490 | - | - | - |
| BAS reward responsivity | 3.41 | 0.38 | 2.40-4.00 | 3.33 | 0.34 | 2.60-4.00 | - | - |  | 2.561 | 0.1110 | - | - | - |
| BAS – BIS | 0.27 | 0.61 | -1.37-2.13 | 0.18 | 0.59 | -1.25-1.58 | - | - |  | 1.077 | 0.3000 | - | - | - |
| Gender |  | | | | | | | |  | χ^2^ | |  | | |
|  |  |  |  |  |  |  |  |  |  | test value | p-value |  |  |  |
|  | 42 women | |  | 73 women | |  | 84 women | |  | 53.636 | **<0.0001** | **<0.0001** | **0.0001** | **<0.0001** |

Table S2 Available data for each of the investigated samples. x marks available datasets.

| Data | Sample 1 (n=117) | Sample 2 (n=89) | Sample 3 (n=152) |
| --- | --- | --- | --- |
| TFEQ | x | x | x |
| BIS/BAS | x | x |  |
| Anthropometric data (BMI) | x | x | x |
| rsEEG | x |  |  |
| rsfMRI | x | x | x |

TFEQ – three factor eating questionnaire; BIS/BAS – behavioural activation/inhibition system questionnaire; rsEEG – resting-state EEG; rsfMRI – resting-state fMRI

Table S3 Table with results of multiple regression analyses investigating the relationship between EEG asymmetry indices and approach/avoidance questionnaire measures for the mean of eyes open and eyes closed conditions. Statistically significant coefficients have been marked in bold. Note that the p-value threshold after Bonferroni correction for four separate regression analyses is 0.0125. RC – rotated component, RR – reward responsivity

|  | Alpha frontal | | Low alpha frontal | | Alpha parietal | | Low alpha parietal | |
| --- | --- | --- | --- | --- | --- | --- | --- | --- |
|  | Beta | p-value | Beta | p-value | Beta | p-value | Beta | p-value |
| BAS fun | -0.50 | 0.0663 | -0.42 | 0.0637 | 0.26 | 0.4027 | 0.37 | 0.5733 |
| BAS fun * gender | 0.21 | 0.3165 | 0.29 | 0.0905 | -0.27 | 0.4419 | -0.31 | 0.6379 |
| BAS drive | -0.10 | 0.7225 | 0.53 | 0.0239 | 0.27 | 0.8627 | -0.02 | 1.0000 |
| BAS drive * gender | -0.01 | 1.0000 | -0.54 | 0.0449 | -0.40 | 0.4815 | 0.20 | 0.3875 |
| BAS RR | 0.45 | 0.0925 | 0.55 | 0.0449 | -0.06 | 0.6429 | -0.19 | 0.3899 |
| BAS RR * gender | -0.49 | 0.0895 | -0.75 | 0.0178 | -0.03 | 0.6604 | -0.01 | 0.9412 |
| BAS - BIS | 0.14 | 1.0000 | -0.52 | 0.2194 | -0.66 | 0.276 | -0.51 | 0.0995 |
| BAS – BIS * gender | -0.01 | 1.0000 | 0.86 | 0.0134 | -0.93 | 0.0132 | 0.44 | 0.1942 |
| Age | -0.05 | 0.3851 | -0.03 | 0.8431 | 0.01 | 1.0000 | -0.12 | 0.1257 |
| BMI | 0.22 | 0.1450 | 0.30 | 0.0164 | -0.25 | 0.0310 | -0.16 | 0.2711 |
| Gender | 0.15 | 0.0663 | 0.37 | 0.0905 | 0.41 | 0.2892 | 0.37 | 0.3474 |

Table S4 Results of multiple regression analyses investigating the relationship between EEG asymmetry indices and eating questionnaire measures for the eyes open condition. Note that the p-value threshold after Bonferroni correction for four separate regression analyses is 0.0125. CC – cognitive control, DI – disinhibition.

|  | Alpha frontal | | Low alpha frontal | | Alpha parietal | | Low alpha parietal | |
| --- | --- | --- | --- | --- | --- | --- | --- | --- |
|  | Beta | p-value | Beta | p-value | Beta | p-value | Beta | p-value |
| CC | -0.11 | 0.5476 | -0.02 | 0.8627 | 0.28 | 0.2444 | 0.26 | 0.5810 |
| CC * gender | 0.11 | 0.5476 | -0.11 | 0.6863 | -0.36 | 0.1537 | -0.16 | 0.7840 |
| DI | 0.26 | 0.5000 | 0.23 | 0.4110 | 0.28 | 0.0966 | 0.12 | 0.6230 |
| DI * gender | -0.27 | 0.4951 | -0.23 | 0.4583 | -0.29 | 0.2210 | -0.16 | 0.6670 |
| Age | 0.01 | 0.9804 | 0.03 | 1.0000 | 0.11 | 0.8039 | -0.02 | 0.9220 |
| BMI | 0.21 | 0.0292 | 0.14 | 0.5217 | -0.14 | 0.6029 | -0.05 | 0.6600 |
| Gender | 0.35 | 0.1625 | 0.45 | 0.0502 | 0.33 | 0.1419 | 0.16 | 0.7650 |

Table S5 Results of multiple regression analyses investigating the relationship between EEG asymmetry indices and eating questionnaire measures for the mean of eyes open and eyes closed condition. Note that the p-value threshold after Bonferroni correction for four separate regression analyses is 0.0125. CC – cognitive control, DI – disinhibition.

|  | Alpha frontal | | Low alpha frontal | | Alpha parietal | | Low alpha parietal | |
| --- | --- | --- | --- | --- | --- | --- | --- | --- |
|  | Beta | p-value | Beta | p-value | Beta | p-value | Beta | p-value |
| CC | -0.02 | 1.0000 | 0.09 | 0.5000 | 0.31 | 0.7255 | 0.30 | 0.1620 |
| CC * gender | 0.01 | 1.0000 | -0.18 | 0.4180 | -0.15 | 0.7451 | -0.27 | 0.2100 |
| DI | 0.07 | 0.9410 | 0.23 | 0.2550 | 0.18 | 0.5811 | 0.14 | 0.5100 |
| DI * gender | -0.01 | 1.0000 | -0.30 | 0.1410 | -0.29 | 0.2207 | -0.21 | 0.2100 |
| Age | -0.02 | 0.6380 | -0.05 | 0.5730 | -0.01 | 1.0000 | -0.13 | 0.3240 |
| BMI | 0.18 | 0.784 | 0.27 | 0.0150 | -0.28 | 0.0215 | -0.10 | 0.5100 |
| Gender | 0.06 | 0.8360 | 0.22 | 0.4180 | 0.39 | 0.1473 | 0.36 | 0.1010 |

Table S6 Results of multiple regression analyses investigating the relationship between fMRI asymmetry indices (Sample 1) and eating behaviour. Note that the p-value threshold after Bonferroni correction for four separate regression analyses is 0.0100. The components have been ordered according to decreasing variance explained. CC – cognitive control; DI – disinhibition, RC – rotated component.

|  | RC2 | | RC1 | | RC4 | | RC5 | | RC3 | |
| --- | --- | --- | --- | --- | --- | --- | --- | --- | --- | --- |
|  | Beta | p-value | Beta | p-value | Beta | p-value | Beta | p-value | Beta | p-value |
| CC | -0.15 | 0.9410 | -0.31 | 0.0802 | -0.09 | 0.5833 | 0.32 | 0.0604 | -0.24 | 0.1940 |
| CC * gender | 0.29 | 0.1960 | -0.02 | 1.0000 | -0.05 | 1.0000 | -0.53 | 0.0201 | 0.30 | 0.1170 |
| DI | -0.03 | 0.8430 | -0.02 | 0.7647 | 0.12 | 0.4265 | -0.13 | 0.1670 | -0.05 | 0.9410 |
| DI * gender | 0.12 | 0.4050 | -0.02 | 1.0000 | -0.16 | 0.3961 | 0.17 | 0.4265 | -0.14 | 0.6550 |
| Age | 0.11 | 0.2010 | 0.06 | 0.3657 | -0.15 | 0.1601 | -0.09 | 1.0000 | 0.06 | 0.5940 |
| BMI | -0.16 | 0.4230 | -0.02 | 0.9804 | -0.17 | 0.8824 | 0.11 | 0.1537 | -0.02 | 0.7060 |
| Gender | -0.09 | 0.7840 | -0.21 | 0.4296 | 0.50 | 0.0386 | 0.07 | 0.4815 | -0.44 | 0.1440 |

Table S7 Component loadings for each of the PCA’s rotated components (Sample 1) in the Three Factor Eating Questionnaire analysis. ROIs represent 9 regions of interest selected for the fMRI analyses. BA – Brodmann area, VTA – ventral tegmental area, NAcc – nucleus accumbens, ParacG – paracentral gyrus, PostcG – postcentral gyrus, ROI – region of interest, RC – rotated component.

| ROI | RC2 | RC1 | RC4 | RC5 | RC3 |
| --- | --- | --- | --- | --- | --- |
| BA10 | 0.60 | 0.09 | 0.17 | 0.52 | -0.20 |
| BA9 | 0.42 | 0.68 | -0.08 | 0.02 | -0.15 |
| BA8 | 0.25 | 0.72 | 0.12 | 0.15 | -0.12 |
| BA46 | 0.77 | 0.16 | -0.02 | -0.04 | 0.06 |
| NAcc | -0.05 | 0.00 | 0.92 | -0.06 | 0.01 |
| VTA | -0.24 | 0.80 | -0.03 | 0.13 | 0.12 |
| BA7 | -0.01 | 0.19 | -0.12 | 0.91 | 0.05 |
| ParacG | 0.67 | -0.04 | -0.49 | 0.05 | 0.00 |
| PostcG | 0.00 | -0.06 | 0.01 | 0.01 | 0.97 |
| Cumulative variance explained | 0.19 | 0.38 | 0.50 | 0.63 | 0.75 |

Table S8 Results of multiple regression analyses investigating the relationship between fMRI asymmetry indices (Sample 2) and approach/avoidance questionnaire measures. Note that the p-value threshold after Bonferroni correction for four separate regression analyses is 0.0100. The components have been ordered according to decreasing variance explained. RC – rotated component, RR – reward responsivity.

|  | RC5 | | RC3 | | RC2 | | RC1 | | RC4 | |
| --- | --- | --- | --- | --- | --- | --- | --- | --- | --- | --- |
|  | Beta | p-value | Beta | p-value | Beta | p-value | Beta | p-value | Beta | p-value |
| BAS fun | -0.10 | 0.3111 | 0.09 | 0.4860 | -0.11 | 0.5050 | -0.04 | 0.7843 | -0.02 | 0.7059 |
| BAS fun * gender | 0.08 | 1.0000 | -0.51 | 0.0715 | -0.45 | 0.1600 | -0.30 | 0.7059 | 0.58 | 0.0332 |
| BAS drive | -0.03 | 0.7843 | -0.29 | 0.0416 | -0.06 | 0.8820 | 0.17 | 0.1433 | -0.11 | 0.6230 |
| BAS drive * gender | 0.29 | 0.3728 | 0.57 | 0.0348 | -0.03 | 0.9220 | -0.44 | 0.0926 | 0.01 | 1.0000 |
| BAS RR | 0.26 | 0.0412 | -0.05 | 0.5942 | 0.20 | 0.1260 | -0.05 | 0.6863 | -0.06 | 0.6667 |
| BAS RR * gender | -0.49 | 0.0317 | 0.34 | 0.1245 | -0.32 | 0.2380 | 0.26 | 0.2927 | -0.13 | 0.6029 |
| BAS - BIS | 0.03 | 0.9608 | 0.11 | 0.6190 | 0.02 | 0.5620 | -0.17 | 0.2797 | -0.14 | 0.6863 |
| BAS – BIS * gender | 0.08 | 0.9020 | -0.35 | 0.2915 | -0.23 | 0.2820 | 0.16 | 0.7843 | -0.20 | 0.3678 |
| Age | 0.26 | 0.0217 | 0.04 | 0.6333 | 0.02 | 0.8820 | 0.11 | 0.5275 | 0.16 | 0.0977 |
| BMI | 0.11 | 0.8431 | -0.07 | 0.5275 | -0.10 | 0.6550 | -0.14 | 0.2551 | 0.08 | 0.8431 |
| Gender | -0.19 | 0.4815 | 0.50 | 0.0692 | 0.55 | 0.1400 | -0.03 | 0.7451 | 0.27 | 0.3165 |

Table S9 Component loadings for each of the PCA’s rotated components (Sample 2) in the BIS/BAS and TFEQ analysis. ROIs represent 9 regions of interest selected for the fMRI analyses. BA – Brodmann area, VTA – ventral tegmental area, NAcc – nucleus accumbens, ParacG – paracentral gyrus, PostcG – postcentral gyrus, ROI – region of interest, RC – rotated component.

| ROI | RC5 | RC3 | RC2 | RC1 | RC4 |
| --- | --- | --- | --- | --- | --- |
| BA10 | 0.02 | 0.15 | -0.01 | 0.86 | -0.16 |
| BA9 | 0.22 | 0.75 | -0.06 | 0.11 | -0.29 |
| BA8 | 0.53 | 0.39 | 0.00 | 0.31 | 0.02 |
| BA46 | 0.11 | -0.27 | -0.61 | 0.55 | -0.05 |
| NAcc | -0.01 | -0.01 | -0.08 | -0.17 | 0.88 |
| VTA | 0.30 | -0.68 | 0.00 | 0.01 | -0.26 |
| BA7 | 0.78 | -0.11 | -0.12 | -0.18 | -0.22 |
| ParacG | 0.06 | -0.14 | 0.90 | 0.03 | -0.08 |
| PostcG | 0.64 | -0.05 | 0.27 | 0.22 | 0.34 |
| Cumulative variance explained | 0.16 | 0.31 | 0.45 | 0.59 | 0.71 |

Table S10 Results of multiple regression analyses investigating the relationship between fMRI asymmetry indices (Sample 2) and eating behaviour. Note that the p-value threshold after Bonferroni correction for four separate regression analyses is 0.0100. The components have been ordered according to decreasing variance explained. CC – cognitive control, DI – disinhibition, RC – rotated component.

|  | RC5 | | RC3 | | RC2 | | RC1 | | RC4 | |
| --- | --- | --- | --- | --- | --- | --- | --- | --- | --- | --- |
|  | Beta | p-value | Beta | p-value | Beta | p-value | Beta | p-value | Beta | p-value |
| CC | 0.02 | 0.9412 | 0.08 | 0.3700 | -0.10 | 0.4350 | 0.16 | 0.4810 | -0.08 | 0.2983 |
| CC * gender | 0.29 | 0.5811 | -0.07 | 1.0000 | 0.31 | 0.9020 | -0.20 | 0.3070 | -0.34 | 0.2444 |
| DI | 0.03 | 0.8431 | 0.01 | 1.0000 | 0.07 | 0.8040 | 0.09 | 0.2890 | 0.00 | 1.0000 |
| DI * gender | 0.46 | 0.1789 | -0.01 | 1.0000 | 0.03 | 0.6550 | -0.02 | 1.0000 | 0.72 | 0.0146 |
| Age | **0.30** | **0.0048** | 0.09 | 0.2020 | -0.01 | 1.0000 | 0.08 | 0.5710 | 0.22 | 0.0499 |
| BMI | 0.09 | 0.2824 | -0.13 | 1.0000 | -0.08 | 0.6060 | -0.17 | 0.1330 | 0.06 | 1.0000 |
| Gender | -0.12 | 0.9020 | 0.26 | 1.0000 | 0.36 | 0.2330 | -0.08 | 0.7840 | -0.12 | 0.4775 |

Table S11 Results of multiple regression analyses investigating the relationship between fMRI asymmetry indices (Sample 3) and eating behaviour. Note that the p-value threshold after Bonferroni correction for four separate regression analyses is 0.0100. The components have been ordered according to decreasing variance explained. CC – cognitive control, DI – disinhibition, RC – rotated component.

|  | RC1 | | RC5 | | RC3 | | RC2 | | RC4 | |
| --- | --- | --- | --- | --- | --- | --- | --- | --- | --- | --- |
|  | Beta | p-value | Beta | p-value | Beta | p-value | Beta | p-value | Beta | p-value |
| CC | -0.08 | 0.7450 | 0.09 | 0.1947 | -0.17 | 0.7451 | 0.18 | 0.1020 | -0.05 | 0.5733 |
| CC * gender | 0.25 | 0.8630 | 0.14 | 0.3125 | 0.20 | 0.2062 | -0.01 | 1.0000 | 0.01 | 0.9608 |
| DI | -0.04 | 1.0000 | 0.10 | 0.5714 | 0.12 | 0.2500 | 0.00 | 0.9410 | 0.02 | 0.7647 |
| DI * gender | -0.12 | 0.4300 | -0.25 | 0.1154 | 0.02 | 0.9804 | -0.09 | 0.8820 | -0.27 | 0.1009 |
| Age | 0.08 | 0.3110 | 0.01 | 0.9608 | 0.20 | 0.0176 | -0.05 | 0.5400 | 0.07 | 0.2880 |
| BMI | 0.02 | 1.0000 | 0.14 | 0.0479 | 0.03 | 0.7451 | -0.11 | 0.6330 | -0.11 | 0.0857 |
| Gender | -0.12 | 1.0000 | 0.18 | 0.4737 | -0.40 | 0.0142 | 0.16 | 0.8820 | 0.01 | 0.9608 |

Table S12 Component loadings for each of the PCA’s rotated components (Sample 3) in the TFEQ analysis. ROIs represent 9 regions of interest selected for the fMRI analyses. BA – Brodmann area, VTA – ventral tegmental area, NAcc – nucleus accumbens, ParacG – paracentral gyrus, PostcG – postcentral gyrus, ROI – region of interest, RC – rotated component.

| ROI | RC1 | RC5 | RC3 | RC2 | RC4 |
| --- | --- | --- | --- | --- | --- |
| BA10 | 0.28 | 0.63 | -0.09 | 0.01 | -0.09 |
| BA9 | 0.78 | 0.07 | 0.02 | 0.06 | -0.04 |
| BA8 | 0.81 | 0.10 | 0.07 | 0.02 | 0.00 |
| BA46 | -0.05 | 0.86 | 0.00 | -0.06 | 0.05 |
| NAcc | 0.05 | -0.08 | -0.04 | 0.91 | -0.05 |
| VTA | -0.04 | -0.03 | -0.02 | -0.03 | 0.96 |
| BA7 | 0.35 | 0.20 | -0.67 | -0.11 | 0.21 |
| ParacG | 0.29 | -0.08 | 0.70 | -0.27 | 0.04 |
| PostcG | 0.17 | 0.29 | 0.54 | 0.39 | 0.22 |
| Cumulative variance explained | 0.18 | 0.32 | 0.46 | 0.58 | 0.70 |
